# Engineering Sustained-Release Broadly Neutralizing Antibody Formulations

**DOI:** 10.1101/2025.05.27.656504

**Authors:** Carolyn K. Jons, Catherine M. Kasse, Bryan T. Mayer, Ollivier Hyrien, Samya Sen, Emily L. Meany, Andrea I. d’Aquino, Priya Ganesh, Noah Eckman, Changxin Dong, Jerry Yan, Leslee T. Nguyen, Vanessa M. Doulames, Ye Eun Song, Olivia M. Saouaf, Christian M. Williams, Shoshana C. Williams, Jesus Paredes, Rajeev Raghavan, Martina Palomares, Michael Alpert, Nicole L. Yates, Georgia D. Tomaras, Michael S. Seaman, Michael Farzan, Eric A. Appel

## Abstract

Sustained serum levels of broadly neutralizing antibodies (bnAbs) are crucial for effective passive immunization against infectious diseases as protection persists only while these bnAbs remain at adequate concentrations within the body. Current obstacles, such as poor pharmacokinetics (PK) and burdensome administration, must be overcome to make bnAbs a viable option for pre- and post-exposure prophylaxis. In this work, we explore how a polymer-nanoparticle (PNP) hydrogel depot technology can be engineered to prolong protein delivery. In-vivo studies in mice and rats demonstrate prolonged protein release, and modeling efforts predict the impact of both the elimination half-life of the active pharmaceutical ingredient and hydrogel depot volume on overall pharmacokinetics. Moreover, flow cytometry characterization reveals that immune cell infiltration into the hydrogel depot can result in faster-than-expected release of antibody cargo on account of active transport via cellular uptake. We then demonstrate that co-formulation of antibodies with an anti-inflammatory agent reduces cellular infiltration and resulting active transport, further extending delivery and pharmacokinetics. Finally, multicompartmental modeling predicts the human PK profiles of clinically relevant HIV bnAbs delivered via subcutaneous hydrogel injection. These findings aid in the development of next generation hydrogel materials that stabilize and slowly release bnAbs for long-term pre-exposure immunoprophylaxis.

## 1. Introduction

Passive immunization, which involves the direct administration of neutralizing antibodies from an external source, offers an essential alternative to vaccination, particularly in situations where an effective vaccine has yet to be developed, or an individual cannot produce their own antibodies (*1–3*). Unlike active immunization, which depends on the body’s immune response and requires time to develop protection, passive immunization provides immediate defense against infections. Passive immunization can occur maternally—through breastfeeding or the placenta—or via intravenous delivery of antibodies (**Fig 1a**) (*4, 5*). Passive immunization is highly relevant to pathogens like HIV, malaria, and Zika, where vaccine development is challenged by high mutation rates and poorly understood immunity markers (*6–12*). In such cases, the identification and administration of broadly neutralizing antibodies (bnAbs) make passive immunization a viable strategy for pre-exposure prophylaxis, post-exposure prophylaxis, and treatment (*2, 5*).

**Figure 1.**
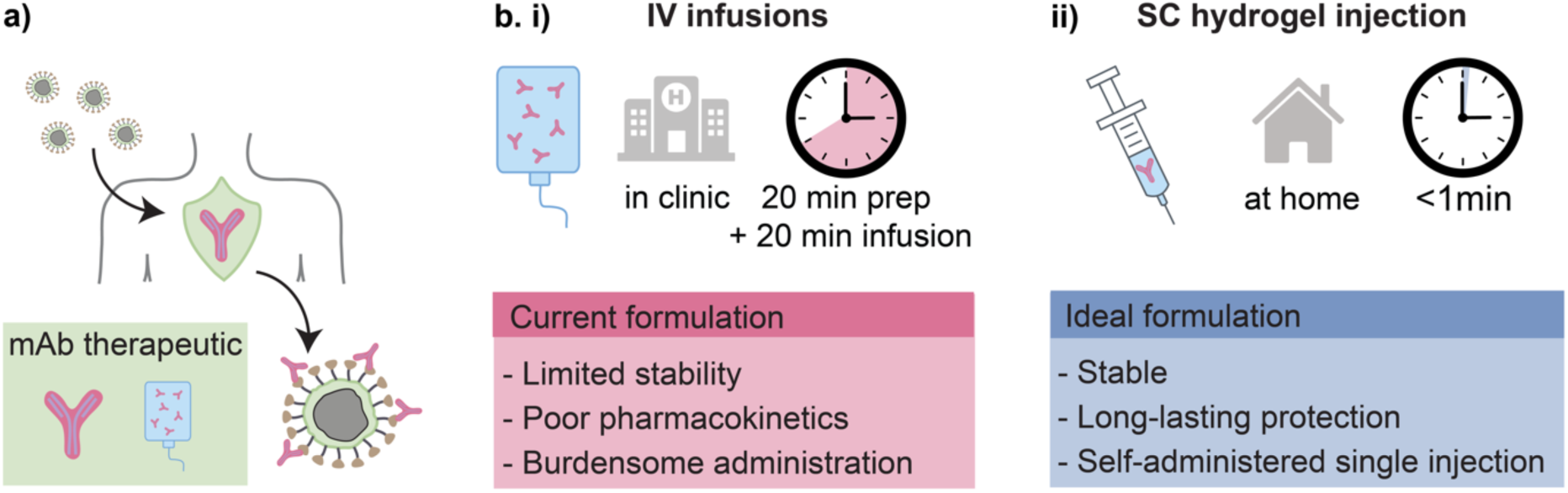
Depot technologies enable extended mAb delivery for passive immunization. **a)** mAb therapeutics in circulation enable neutralization of virus. **b)** mAb therapeutics are typically administered in hospital settings via **(i)** intravenous infusion. An ideal **(ii)** subcutaneous hydrogel delivery technology would provide extended mAb release following a single, at-home injection.

While passive immunization is an exciting strategy to combat infectious disease, it has several key limitations. First, although passive immunization provides almost immediate protection, an individual is only protected for as long as antibodies remain circulating in the blood stream at therapeutically relevant concentrations. Many bnAbs exhibit short circulatory half-lives (days to weeks) which heavily limits their utility for long term pre-exposure prophylaxis (*13*). Additionally, the necessity of repeat doses to maintain therapeutic efficacy reduces adherence and increases the burden on healthcare systems. Because monoclonal antibodies are often delivered intravenously, their administration is time-consuming and generally restricted to clinical settings, making administration on a population-wide basis or in resource-limited environments challenging (**Fig. 1b i**) (*14*).

To make passive immunization more practical, a key objective is to extend the retention time of the bnAbs in the body (*15*). One effective approach is to engineer antibodies for extended half-life, particularly by modifying their crystallizable fragment (Fc) region, which has been proven to significantly prolong their presence in the bloodstream (*16–19*). Advancements in bnAb engineering have largely stemmed from passive HIV prophylaxis research with several long-acting bnAbs incorporating Fc modifications now undergoing clinical trials (*12, 16, 20–22*). While Fc modification extends circulatory half-life, antibody engineering is not a viable strategy for all antibodies as Fc modification can negatively impact antibody function, increase immunogenicity, and heighten manufacturing and regulatory challenges (*23*).

Another promising and complementary approach to improving bnAb circulation time is the development of materials that can act as sustained-release depots (**Fig. 1b ii**) (*24–26*). Depot technologies extend the time biologics remain in blood circulation by delaying release from the subcutaneous space into the bloodstream. Hydrogels, which are cross-linked, water-swollen macromolecular networks, are a promising materials class for use as drug depots (*27–35*). Their high water content provides physicochemical similarity to biological tissues and enables encapsulation of hydrophilic drug cargo, while the presence of a polymer network provides physical structure, tunable mechanical properties, and controlled cargo diffusivity (*36*). In particular, physically crosslinked hydrogels are a promising materials class for bnAb delivery as they exhibit shear-thinning and self-healing behaviors which enable facile injectability and depot formation upon administration (*36–41*). These materials can be administered via standard needles or auto injectors enabling a minimally invasive strategy to prolong delivery of antibody therapeutics (*42, 43*). However, despite these demonstrated advantages, the design of hydrogel depots for long-term antibody delivery remains challenging. These include engineering networks with sufficiently low cargo diffusivity for sustained release, ensuring a high degree of hydrogel biocompatibility over extended residence times, and understanding immune responses to antibody-loaded materials within the subcutaneous space.

In this study, we aim to develop broader insights into the design of hydrogel-based antibody depots for extended delivery of bnAbs. Using in-vivo pharmacokinetics (PK) studies and compartmental modeling we assess how antibody half-life and depot volume influence release kinetics and pharmacokinetics. We also examine how immune cell infiltration into the hydrogel depot impacts cargo release over time. These findings support the development of next-generation hydrogel-based bNAb drug product candidates for long-term immunoprophylaxis.

## 2. Results and Discussion

### 2.1 Rheological and transport properties of PNP hydrogels

The Appel lab has previously reported the use of polymer-nanoparticle (PNP) hydrogels for sustained biotherapeutic delivery (*43–49*). PNP hydrogels are physically crosslinked hydrogels formed through dynamic, multivalent interactions between hydrophobically modified hydroxypropylmethylcellulose (HPMC-C_12_) and nanoparticles prepared via nanoprecipitation of poly(ethylene glycol)-block-poly(lactic acid) (PEG-PLA NPs) (**Fig. 2a**). Unlike conventional uses of these nanoparticles as nanocarriers for drug delivery applications, in PNP hydrogels they instead function as structural crosslinking nodes that physically link HPMC-C_12_ chains to form a percolated network with a small mesh size able to entrap biologic cargo (*50–52*). These materials are easily formulated by mixing aqueous solutions of the two structural components allowing for mild gelation conditions suitable for encapsulating biotherapeutic cargo such as antibodies (*48*). The mechanical properties of PNP hydrogels can be tuned by adjusting the polymer to nanoparticle ratio as well as the total solids content. In this work, a formulation comprising 2 wt% HPMC-C_12_ and 10 wt% PEG-PLA nanoparticles, with the remaining 88 wt% comprising buffer and drug cargo, will be denoted PNP-2-10. Due to their entropically-driven physical crosslinks (*51*), these hydrogels maintain consistent mechanical properties across relevant temperatures, exhibiting comparable stiffness at storage, injection, and body temperatures (**Fig. 2b**). Furthermore, PNP hydrogels enable high levels of cargo loading, with stiffness remaining unchanged upon incorporation of model protein cargo, human IgG (hIgG) (**Fig. 2c**).

**Figure 2.**
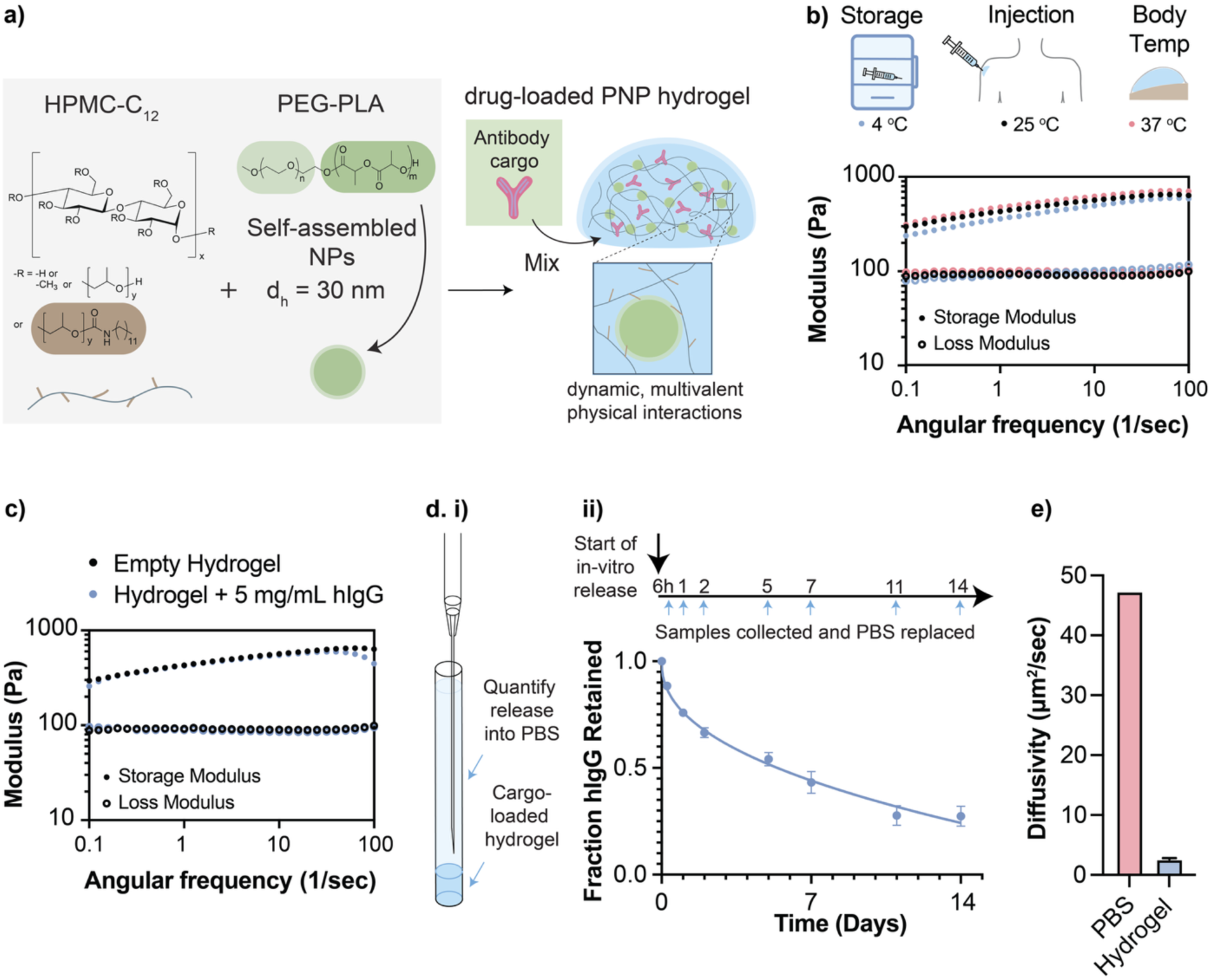
PNP hydrogels enable controlled antibody delivery. **a)** PNP hydrogels are formed from dynamic crosslinking of HPMC-C_12_ and PEG-PLA nanoparticles. Antibody cargo is easily encapsulated during the mixing process. **b)** Frequency sweeps of PNP-2-10 hydrogels at 4 °C, 25 °C, and 37 °C demonstrating temperature invariant rheological properties at storage, injection, and body temperature. **c)** Frequency sweep of empty and antibody loaded (5 mg/mL hIgG) PNP-2-10 hydrogels. **d. i)** Diagram of a capillary release assay for evaluating antibody release from PNP hydrogels into PBS. **ii)** Sample collection schedule (n=4) and corresponding retention of hIgG in the PNP-2-10 hydrogel. **e)** Diffusivity of hIgG in PBS as predicted by Stokes-Einstein law and diffusivity of hIgG in the PNP-2-10 hydrogel as measured by FRAP (n=3). Plots show mean ± SD.

PNP hydrogels represent a promising depot technology for sustained biotherapeutic delivery due to their ability to significantly slow the diffusion of biologic cargo. In vitro capillary release assays demonstrate controlled release of hIgG over a 14-day period (**Fig. 2d**). Release data fit to the Korsmeyer-Peppas model (*53*) demonstrates release from hydrogels is governed by Fickian diffusion (**SI Fig 1**). Fluorescence recovery after photobleaching (FRAP) measurements show that the PNP-2-10 formulation exhibits a low cargo diffusivity of 2.6 µm²/sec (**Fig. 2e**), which is an 18-fold reduction in the diffusivity predicted for hIgG in PBS (**Fig. 2e**) (*54*). Prior work from the Appel lab has also shown that PNP-2-10 hydrogels stabilize biologic cargo by preventing aggregation at ambient temperature (*48, 55*) and, following injection, form a persistent hydrogel depot in the subcutaneous space of mice that lasts for several weeks (*40, 41*).

### 2.2 Pharmacokinetics in mice and compartmental modeling fundamentals

Sustained-release strategies are especially valuable for HIV therapies, where long-term antibody exposure is critical for effective prophylaxis (*2*). A variety of antibodies that bind HIV have been isolated or engineered for use in HIV prevention or treatment (*3*). Two promising HIV bnAbs are eCD4-Ig and PGT121. eCD4-Ig is an engineered antibody-like fusion protein consisting of an extended version of CD4, the primary receptor for HIV entry, fused to an IgG Fc domain. This fusion allows eCD4-Ig to mimic the natural binding of CD4 to the HIV envelope glycoprotein (gp120), preventing viral entry into host cells (*56, 57*). PGT121 targets the HIV-1 envelope glycoprotein (Env), specifically binding to a conserved epitope in the V3 glycan region of gp120 (*22, 58*). eCD4-Ig and PGT121 have elimination half-lives in mice of 5 days and 12 days respectively (**SI Fig 2)**.

To assess the efficacy of PNP hydrogels as a subcutaneous bnAb delivery depot for HIV passive immunization, both standard aqueous formulations and PNP hydrogel-based formulations of eCD4-Ig and PGT121 were prepared. The PK of these formulations were evaluated following subcutaneous administration in scid FcRn-/- hFcRn (32) Tg mice. For eCD4-Ig evaluation, mice received a 360 µg dose administered subcutaneously in either a bolus of a standard vehicle or in 200 µL of PNP-2-10 hydrogel (**Fig. 3a**). For PGT121 evaluation, mice received a 1 mg dose administered subcutaneously either in a bolus of a standard vehicle or in 200 µL of PNP-2-10 hydrogel (**Fig. 3b**). Blood samples were collected over time, and protein concentrations in serum were analyzed by ELISA.

**Figure 3.**
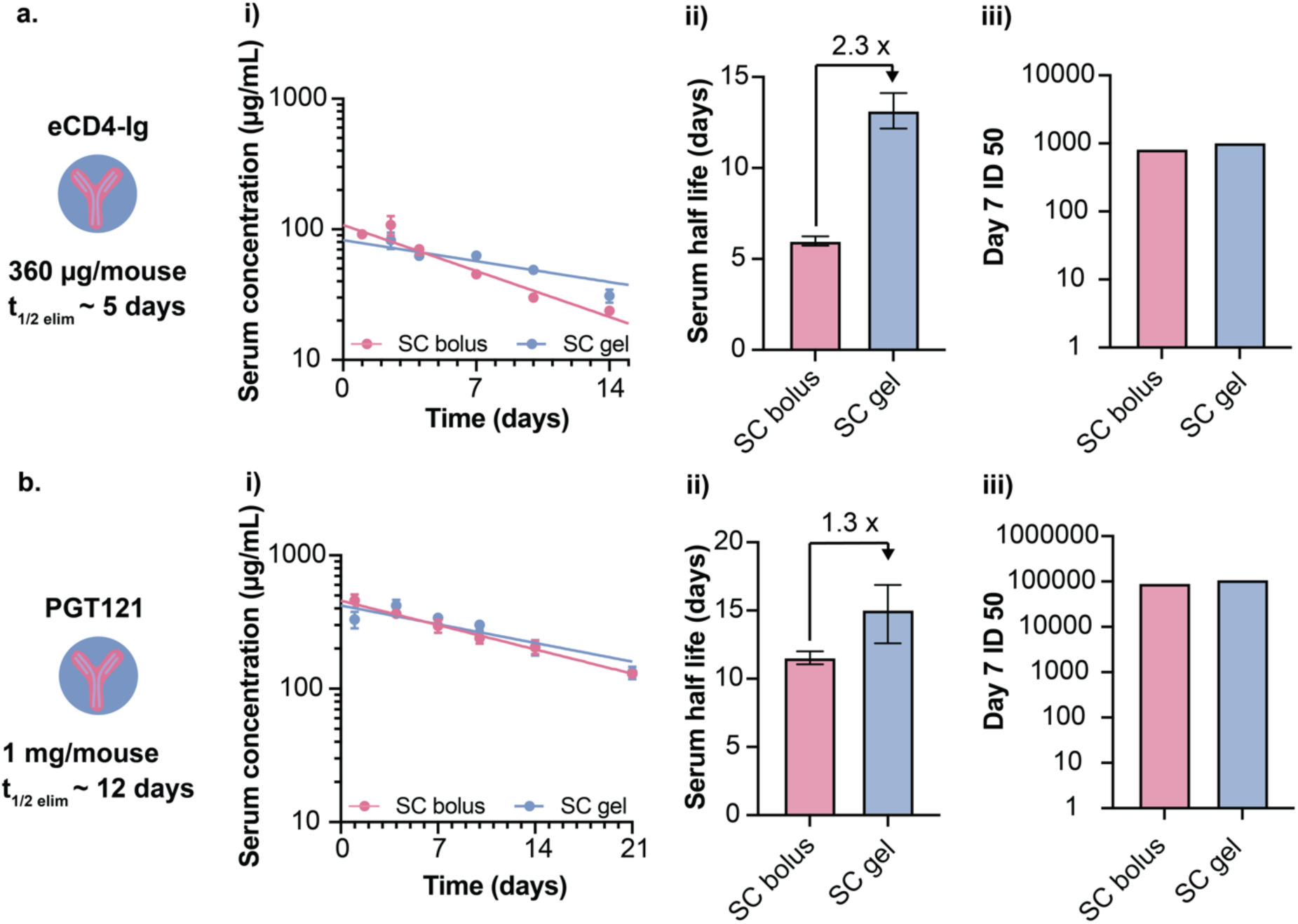
mAb elimination half-life relative to SC absorption half-life impacts degree of observable depot effect. **a)** eCD4-Ig antibody is administered to *scid* FcRn-/- hFcRn mice via a SC bolus or SC gel injection, and PK is assessed via ELISA. **(i)** eCD4-Ig serum concentration profiles from day 2.5 onwards are fit to a single-phase exponential decay to identify **(ii)** serum elimination half-life. Bar graph shows mean ± parameter fit uncertainty. **(iii)** Neutralizing activity of eCD4-Ig from pooled mouse serum (n=4) 7 days post-administration is quantified by TZM-bl neutralization assay. **b)** PGT121 antibody is administered to *scid* FcRn-/- hFcRn mice via a SC bolus or SC gel injection, and PK is assessed via ELISA. **(i)** PGT121 serum concentration profiles from day 4 onwards are fit to a single-phase exponential decay to identify **(ii)** serum elimination half-life. Bar graph shows mean ± parameter fit uncertainty. **(iii)** Neutralizing activity of PGT121 from pooled mouse serum (n=7) 7 days post-administration is quantified by TZM-bl neutralization assay.

A single-phase exponential decay was fit to the resultant serum concentration profiles from day 2.5 onwards for eCD4-Ig and day 4 onwards for PGT121 to identify comparative circulation half-life between treatments (**Fig. 3a i**, **Fig. 3b i, SI Fig 2**). The circulation half-life of eCD4-Ig showed a 2.3-fold improvement when delivered via hydrogel (13 days) as compared to bolus (6 days) (**Fig. 3a ii, SI Fig 3**). By contrast, the circulation half-life of PGT121 showed only a 1.3-fold improvement when delivered via hydrogel (14 days) as compared to bolus (11 days) (**Fig. 3b ii, SI Fig 4**). In both cases, the antibodies retained their neutralization capabilities, exhibiting comparable ID50 values at day 7 between bolus and hydrogel delivery (**Fig. 3a iii**, **Fig. 3b iii**).

PK of subcutaneous bnAb delivery can be described using compartmental modeling (**Fig. 4a, Supplementary Discussion 1**) (*48, 59*). The observed circulation half-life of a mAb is dependent on the combined rate of release from the hydrogel depot and subsequent absorption through the subcutaneous space (t_½ abs_) as well as the elimination half-life of the biologic (t_½ elim_). As bnAbs with shorter half-lives are more rapidly cleared from the serum, we hypothesized the impact of sustained release from a subcutaneous depot on circulation half-life would be more evident. Compartmental modeling with a fixed rate of release from a subcutaneous depot (t_½ abs_) but varied bnAb elimination half-life (t_½ elim_) (**Fig 4b**) illustrated how the relative improvement compared to bolus in time above a therapeutic threshold is greater when mAb elimination half-life is short compared to depot release half-life (**Fig. 4b iii**). These results were illustrated experimentally with hydrogel depots showing greater improvements in bnAb circulation half-life for eCD4-Ig than for PGT121.

**Figure 4.**
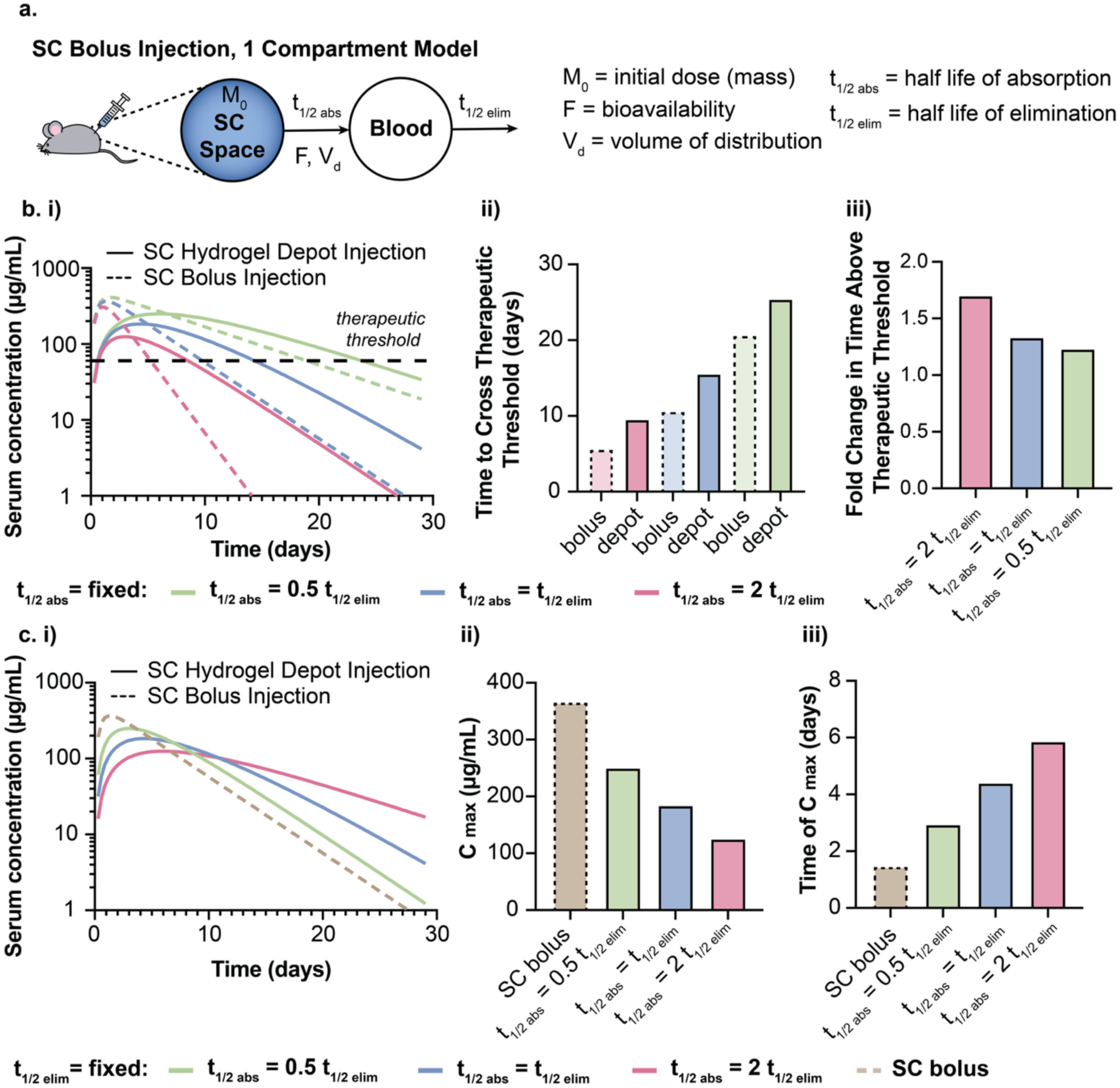
Effects of absorption half-life and elimination half-life on compartment modeling of mAb serum concentration. **a)** One-compartment PK models that include relevant physiological parameters and antibody drug characteristics can be used to model mAb serum concentration following antibody administration via SC bolus or SC gel. **b. i)** Modeled bnAb serum concentration profiles for hydrogel injections with fixed half-lives of SC absorption but varied mAb elimination half-life. Corresponding **ii)** time to cross therapeutic threshold and **iii)** fold change in time above therapeutic threshold for hydrogel administration compared to bolus administration. **c. i)** Modeled bnAb serum concentration profiles for hydrogel depot injections with fixed mAb elimination half-life but varied half-lives of SC absorption. Corresponding serum concentration **ii)** C_max_ and **iii)** time of C_max_.

As antibody engineering efforts continue to yield development of bnAbs with half-life extensions, it is relevant to consider the half-life of cargo release from a depot to effectively engineer a long-acting drug product candidate. Further compartment modeling with fixed bnAb elimination half-life (t_½ elim_) but varied rates of subcutaneous depot release (t_½ abs_) demonstrated that the circulation time of a bnAb can be extended by increasing t_½ abs_ (**Fig. 4c i**). Moreover, an extended bnAb release timeline with an equal initial dose resulted in a delayed and reduced C_max_ (**Fig. 4c ii,iii**). For applications like cancer immunotherapy where dose-dependent toxicity is of concern, delivery strategies that reduce C_max_ improve patient safety (*60*). While high doses of bnAbs are generally not a toxicity concern, reducing C_max_ enables a longer window of therapeutic efficacy for a given dose of drug.

### 2.3 Effect of hydrogel volume of administration on bnAb PK

One approach to increase t_½ abs_ to extend the circulation of a bnAb is to increase the volume of hydrogel administration. Increasing hydrogel volume results in an increase in diffusion length and therefore slower release of entrapped bnAb from the hydrogel depot into the subcutaneous space. While the mouse subcutaneous space is limited to an administration volume of 200 µL, the human subcutaneous space can accommodate much larger volumes of up to 2 mL (*61*).

To assess the impact of larger hydrogel administration volumes on the rate of bnAb release and resultant PK, we used Sprague Dawley-Rag2^em2hera^ll2rg^em1hera^/HblCrl (SRG) rats. These animals received a 2.3 mg dose of PGT121 either IV or SC in a bolus of a standard vehicle, or SC in PNP-2-10 hydrogel injections of 250 µL, 500 µL, or 1 mL. Blood samples were collected over time, and protein concentrations in serum were analyzed by ELISA (**Fig. 5a**). The resultant PGT121 serum concentration profiles were fit to a one-compartment model to obtain half-life of subcutaneous depot release (t_½ abs_) as well as C_max_ and time to C_max_ (**Fig. 5b**). One-compartment model fits utilized t_½ elim_ values fit from IV bolus data (**SI Fig 5**).

**Figure 5.**
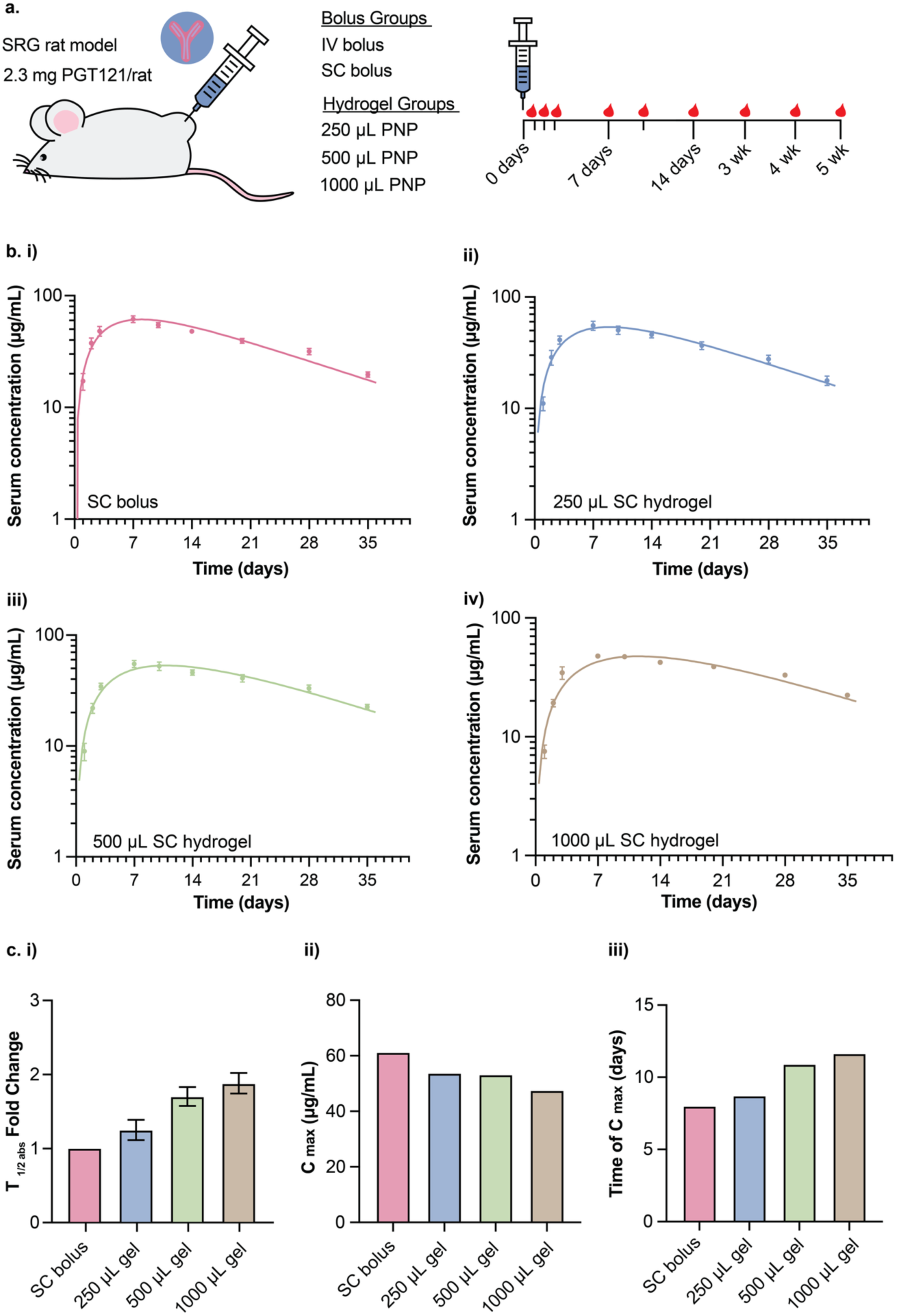
Hydrogel administration volume impacts kinetics of mAb release from the subcutaneous space. **a)** PGT121 antibody is administered to *SRG rats* via a SC bolus or 250 µL, 500 µL, or 1 mL SC gel injection, and PK is assessed via ELISA. **b)** PGT121 serum concentration profiles for **i)** SC bolus, **ii)** 250 µL gel, **iii)** 500 µL gel, or **iv)** 1 mL gel fit to a one-compartment model. **c)** Corresponding **i)** fold change in subcutaneous absorption half-life compared to SC bolus, **ii)** C_max_, and **iii)** time of C_max_ from model fits. Bar graph shows mean ± SEM.

Increasing hydrogel volume resulted in delayed cargo release and an increase in t_½ abs_ (**SI Fig 6**) with 250 µL, 500 µL, and 1 mL hydrogels showing a fold-improvement of 1.2, 1.7, 1.9 over t_½ abs_ of the SC bolus (**Fig. 5c i**). Additionally, one-compartment modeling of PGT121 delivered from larger hydrogel volumes shows an expected decrease in C_max_ as well as a later time to C_max_ (**Fig. 5c ii,iii**). These findings suggest that in addition to designing hydrogel materials with low diffusivities, it is relevant to consider the impact of hydrogel administration volume on cargo release kinetics and PK.

### 2.4 Role of Cellular Infiltration on PK

Many subcutaneously administered biomaterials not only enable controlled cargo release but also interact with the host immune system (*36, 62*). PNP hydrogels facilitate cargo release through both passive diffusion and erosional mechanisms (*63*), while their dynamic crosslinks can enable cellular infiltration and motility (**Fig. 6a i**) (*46, 47*). Cellular infiltration into the hydrogel network is advantageous for vaccine applications, where cellular interactions with vaccine cargo result in more robust immune responses (*47, 49, 64, 65*). However, for passive immunization applications, infiltrating cells may result in restructuring of the hydrogel network and active transport of cargo out of the depot, leading to faster than anticipated therapeutic cargo release.

**Figure 6.**
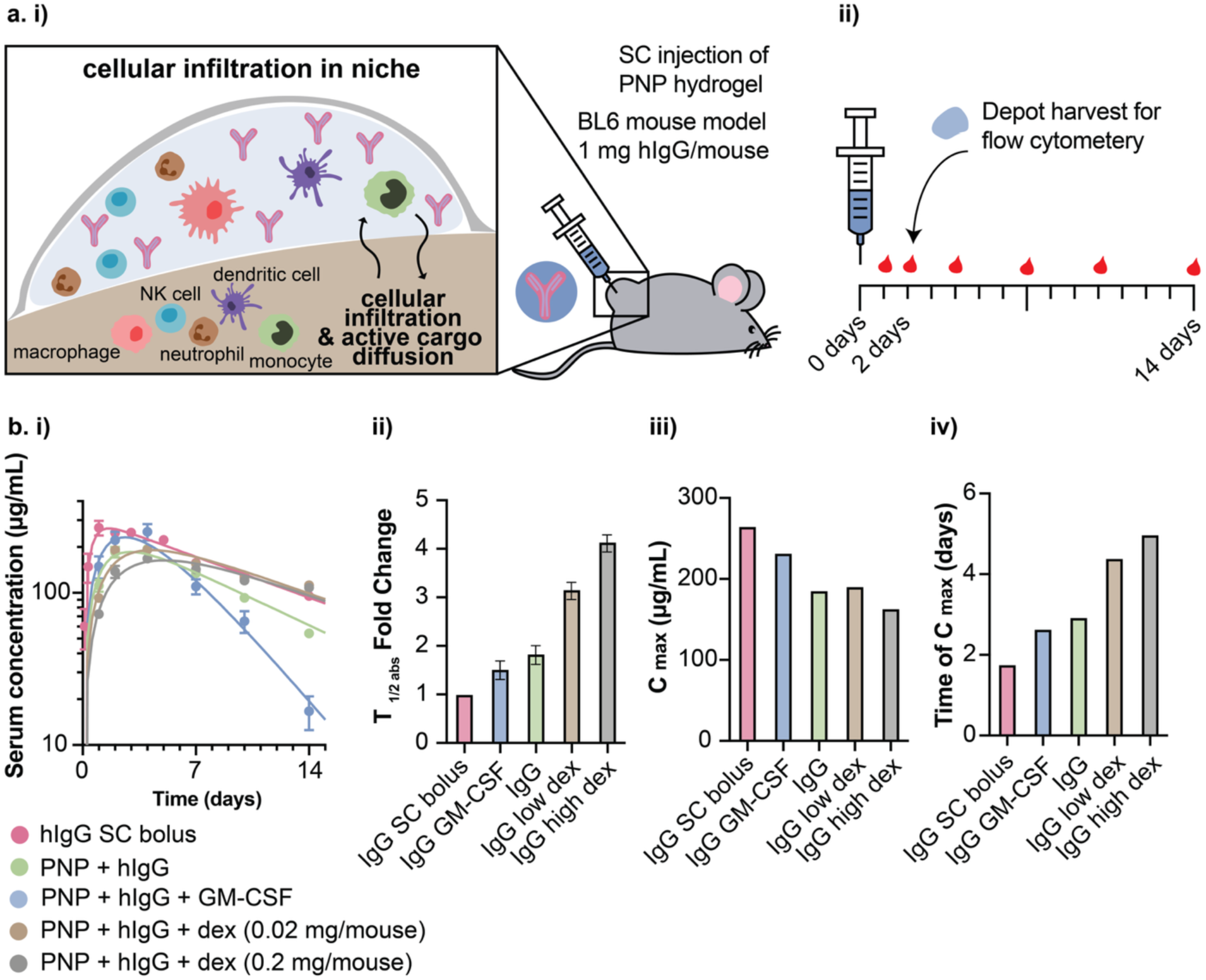
Incorporation of immunomodulatory cargo into hydrogels effects active transport-driven premature mAb release. **a. i)** Dynamic hydrogels allow for cellular infiltration and active cargo diffusion. **(ii)** BL6 mice receive 200 µL SC injections of hydrogels containing empty gel, human IgG (hIgG), hIgG with GM-CSF, hIgG with low dose dexamethasone, or hIgG with high dose dexamethasone. PK is assessed via ELISA. Hydrogel infiltration is assessed in a subset of animals on day 2. **b. i)** hIgG serum concentration profiles are fit to a one-compartment model, with corresponding **ii)** fold change in subcutaneous absorption half-life compared to SC bolus, **iii)** C_max_, and **iv)** time of C_max_ from model fits. Bar graph shows mean ± SEM.

To study the role infiltrating immune cells may play on cargo release, PNP-2-10 hydrogels were prepared with human Immunoglobulin G (hIgG) alone as well as hIgG co-delivered with either immunosuppressant or immunostimulatory cargo. Dexamethasone (dex), a synthetic corticosteroid, was selected as an anti-inflammatory and immunosuppressant cargo (*66*). Granulocyte-Macrophage Colony-Stimulating Factor (GM-CSF) was selected as an immunostimulatory cytokine that attracts immune cells, including phagocytic monocytes and macrophages, and promotes proliferation and activation (*67*). C57BL6 (BL6) mice received a 1 mg dose of hIgG either in a SC bolus or a 200 µL hydrogel injection. Hydrogels contained either hIgG alone, hIgG and GM-CSF, or hIgG and dex at both a low and high dose. Blood samples were collected over time, and hIgG concentrations in serum were analyzed by ELISA (**Fig. 6a ii**). The resultant hIgG PK profiles were fit to a one-compartment model (**Fig. 6b i**) to obtain half-life of subcutaneous depot release (t_½ abs_) as well as C_max_ and time of C_max_ (**Fig. 6b ii,iii,iv**).

The addition of dex resulted in slower cargo release as indicated by an increase in t_½ abs_ (**SI Fig 7**). Hydrogels containing hIgG showed a 1.8-fold increase in t_½ abs_ compared to the SC bolus, while co-formulation with dex at low and high doses resulted in a t_½ abs_ fold increase of 3.2 and 4.1 over bolus respectively (**Fig. 6b ii**). By contrast, co-delivery of GM-CSF in the hydrogel resulted in a decrease in t_½ abs_ compared to the hydrogel with IgG alone (**Fig. 6b ii**). Compartment model fits of C_max_ and time of C_max_ further illustrated hydrogels with higher t_½ abs_ have a lower C_max_ and a later time to C_max_ (**Fig 6b iii,iv**). Except in the case of gels co-delivering GM-CSF, one-compartment model fits were performed using an elimination half-life (t_½ elim_) of 7.85 days, as determined from SC bolus data (**Fig. 6b i**). However, the PK from the co-formulation of IgG with GM-CSF was poorly fit using this same elimination half-life. When t_½ elim_ was allowed to vary, model fitting yielded a shorter elimination half-life of approximately 2 days (**SI Fig 8**). We hypothesize that this decrease in elimination half-life in the presence of GM-CSF may be due to either an increased anti-drug antibody (ADA) response or altered FcRn expression levels or trafficking patterns for immune activation.

To further characterize immune cell infiltrate, 200 µL hydrogels loaded with hIgG, hIgG and dex, or hIgG and GM-CSF were injected into the subcutaneous space of BL6 mice and were excised at day 2 for dissociation and analysis of cellular infiltration with flow cytometry (**Fig. 6a ii, SI Fig 9**). Excised hydrogel depots displayed noticeable differences in cellular infiltration. Hydrogels loaded with hIgG or hIgG and GM-CSF exhibited yellow-tinted cell infiltrate, while those containing hIgG and dex appeared structurally robust and uncolored (**Fig. 7a**). Flow cytometry further quantified cellular infiltration, with hydrogels containing GM-CSF showing an increase in total cells per gel (4.9×10^6^) and hydrogels containing dex showing a decrease in total cells per gel (0.4×10^6^) compared to hydrogels with hIgG alone (0.88×10^6^). The majority of cells found within the hydrogel depots were CD45^+^ leucocytes. (**Fig. 7b i**). Further characterization of the CD45^+^ cell population within the hydrogel niche demonstrated that dex consistently reduced cell counts while GM-CSF consistently increased cell counts uniformly across numerous immune cell types, including myeloid, B, and T cells (**Fig 7b ii**). Flow cytometry analysis of the cellular infiltrate in the hydrogel depot combined with the PK data suggests that incorporating immunosuppressant cargo, such as dex, may prevent cellular infiltration and active cargo diffusion, thereby slowing cargo release from the hydrogel matrix and prolonging PK.

**Figure 7.**
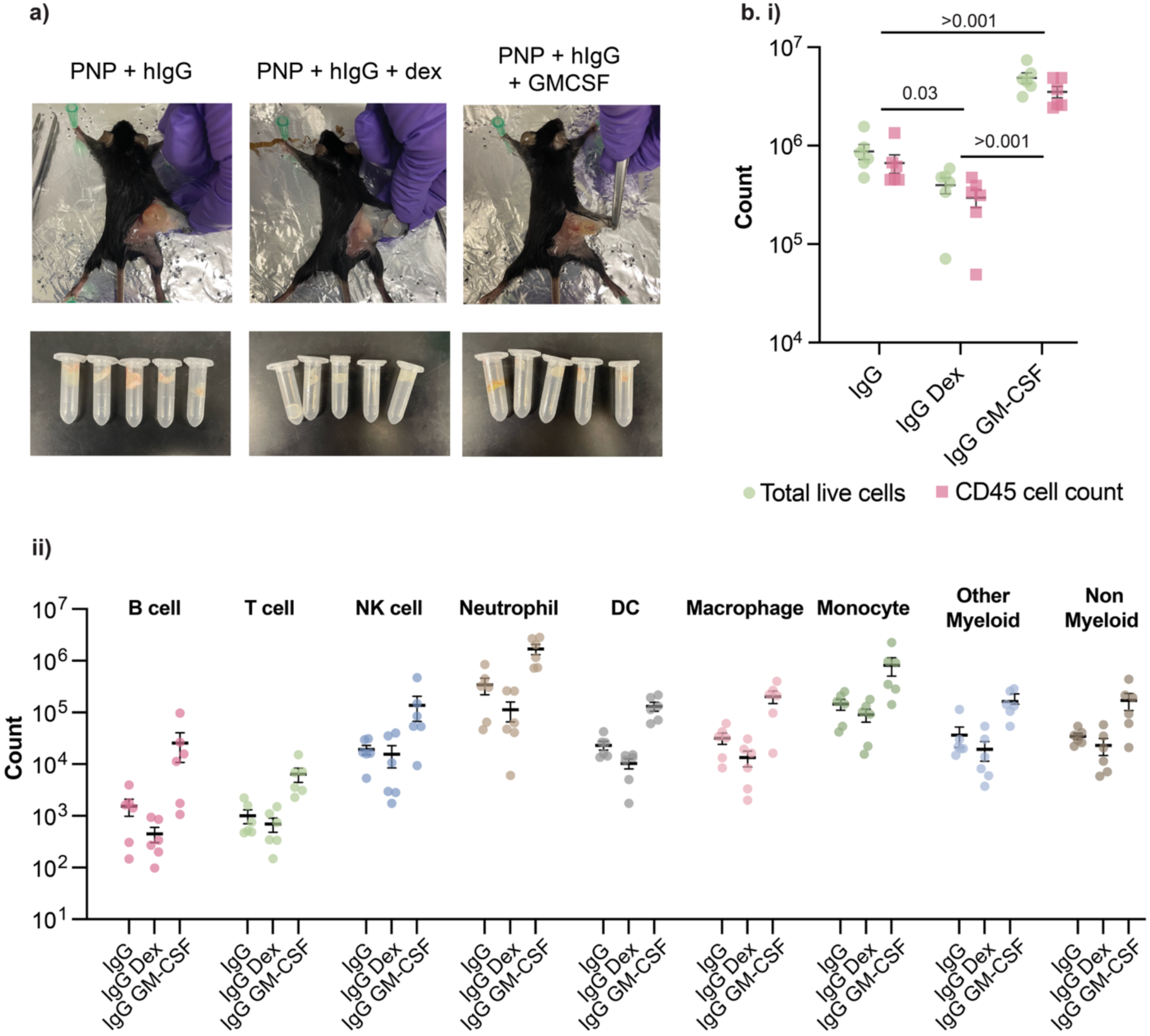
Incorporation of immunomodulatory cargo into hydrogels effects degree of cellular infiltration and infiltrate make up. **a)** Hydrogels loaded with hIgG, hIgG and dex, or hIgG and GM-CSF were injected subcutaneously and excised at day 2 for dissociation and analysis of cellular infiltration with flow cytometry. **b. i)** Total cell counts and counts of CD45+ leukocytes per gel. **ii)** Quantification of cell types in each hydrogel niche. Data shown as mean ± SEM, n = 6. Bar graph shows mean ± SEM. P values from 1-way ANOVA of logged means with Tukey correction for multiple comparisons.

To assess the impact of co-delivery of dex on PGT121 release from a large-volume hydrogel depot, SRG rats received a 1 mL subcutaneous hydrogel injection containing 2.3 mg of PGT121 and 0.5 mg of dex. Blood samples were collected over time, and protein concentrations in serum were analyzed by ELISA (**Fig. 8a**). The resultant PGT121 serum concentration profile (**Fig. 8b**) was fit to one-compartment model to obtain half-life of subcutaneous depot release (t_½ abs_) as well as C_max_ and time of C_max_ (**Fig. 8c**).

**Figure 8.**
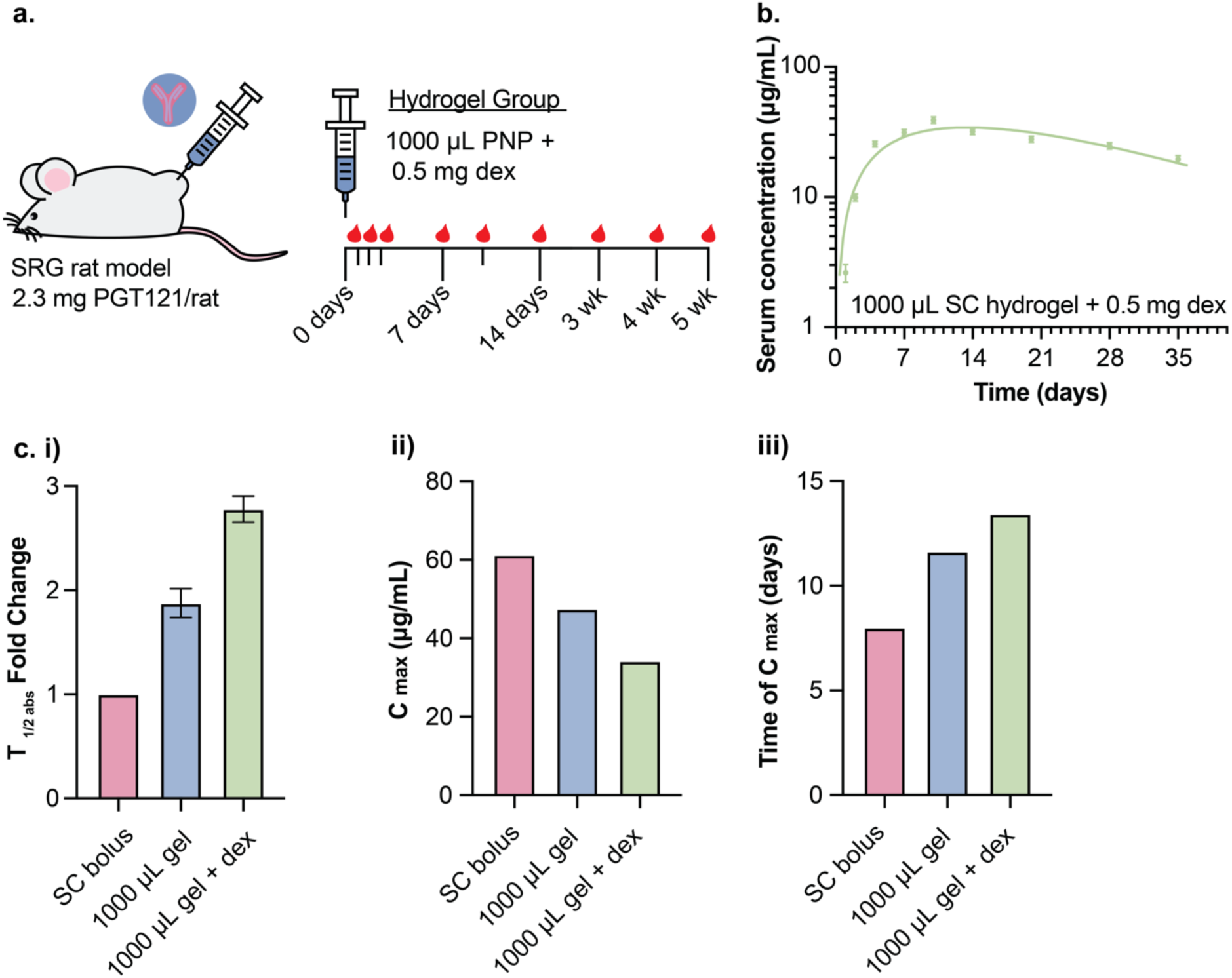
Incorporation of dexamethasone extends mAb release from hydrogels in the subcutaneous space. **a)** PGT121 antibody and dexamethasone are co-administered to *SRG rats* via a 1 mL SC gel injections, and PK is assessed via ELISA. **b)** PGT121 serum concentration from 1 mL gel with dexamethasone is fit to one-compartment model that utilizes elimination half-life fit from IV bolus data. **c)** Corresponding **i)** fold change in absorption half-life compared to SC bolus, **ii)** Cmax, and **iii)** time of Cmax from model fits. Bar graph shows mean ± SEM.

The addition of dex resulted in delayed cargo release and an increase in t_½ abs_ (**SI Fig 10**). Hydrogels containing hIgG and dex showed a fold-improvement of 2.8 over t_½ abs_ of the SC bolus and a fold-improvement of 1.5 over t_½ abs_ of SC gels containing PGT121 alone (**Fig 8c i**). As expected from their reduced diffusivity, hydrogels containing dex further show a decrease in C_max_ and a later time to C_max_ (**Fig. 8c ii,iii**). These findings highlight how the incorporation of immunosuppressant cargo into a hydrogel depot technology can limit active cellular transport of cargo and extend the delivery timeline of bnAbs such as PGT121. While this phenomenon has been clearly observed in PNP hydrogels, we anticipate that incorporating dex into other hydrogel depot systems—particularly those influenced by cellular remodeling or active cellular transport— would similarly result in a slower cargo release rate.

**Figure 10.**
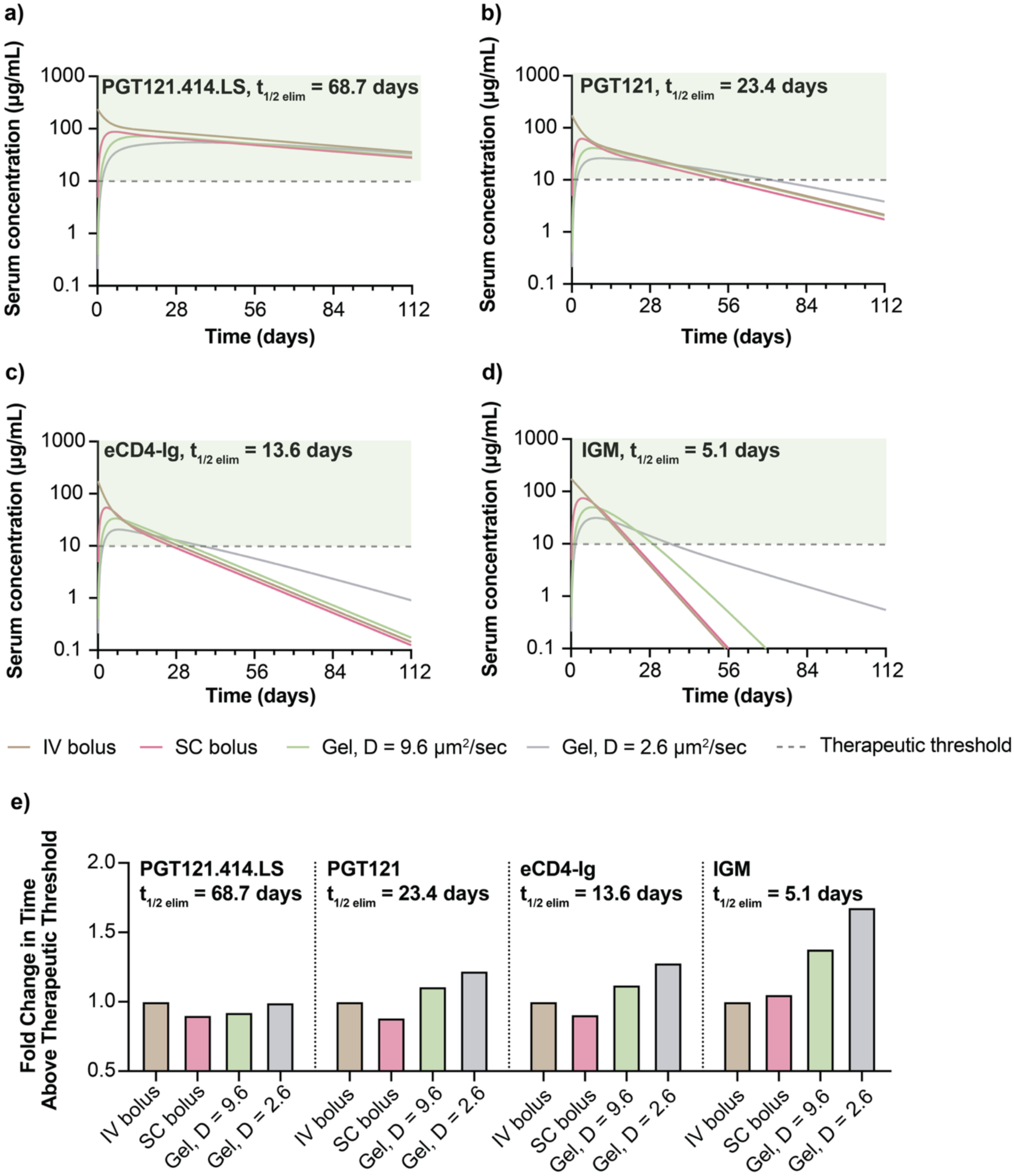
Clinical PK simulations for hydrogel delivery. Modeled protein serum concentration for **a)** PGT121.4414.LS, **b)** PGT121, **c)** eCD4-Ig, and **d)** IGM for IV injection, SC injection, injection of a 2 mL hydrogel with dexamethasone (D = 9.6 µm²/s), and injection of a 2 mL hydrogel with an ideal diffusion coefficient of 2.6 µm²/s. Therapeutic threshold is set at 10 µg/mL. **e)** Fold change in time above therapeutic threshold for subcutaneous bolus or hydrogel administration compared to IV bolus injection.

### 2.5 Multi-compartmental PK and diffusion modeling, and extrapolation to human scales

Thus far, a linear absorption process was used to assess relative rates of antibody release from the subcutaneous space. A multi-step absorption process, however, offers additional detail on PK and depot diffusion properties (*48, 59*). To further explore these dynamics, PGT121 serum concentration profiles for rats receiving PGT121 via either IV bolus, SC bolus, 1 mL hydrogel, or 1 mL hydrogel with dex were fit to a multi-compartment population model accounting for additional PK properties (**Fig. 9a,b**, **Table 1, Supplementary Discussion 2**). First, a two-compartmental PK model with linear absorption was fit to the PGT121 serum concentrations measured in the IV and SC groups (**Fig. 9a**). The model was then extended to include a diffusion component in the SC compartment and was fit to data from the IV, SC, and hydrogel groups (**Fig. 9b**). Incorporating the hydrogel group did not substantially alter key PK parameters, and the model yielded a good fit across all groups, with comparable terminal kinetics (**Table 1**). As observed in one-compartment modeling, addition of dex resulted in a decrease in C_max_ and a later time to C_max_ (**Fig. 9c i,ii**) Notably, while heavily reduced bioavailability is a common limitation of drug delivery technologies, PNP hydrogels showed only a slight reduction in bioavailability comparable to SC bolus dosing (**Fig. 9c iii**). The diffusion coefficient for PGT121 estimated from the model was 27 µm²/s for hydrogels with bnAb alone and 9.6 µm²/s for those comprising bnAb and dexamethasone (**Fig. 9c iv**), indicating that reducing immune cell infiltration decreases the effective diffusion rate of cargo from the hydrogel depot by nearly 3-fold. These results improve our understanding of cargo release kinetics from hydrogel depots in-vivo and can potentially help guide expectations relevant for clinical translation.

**Figure 9.**
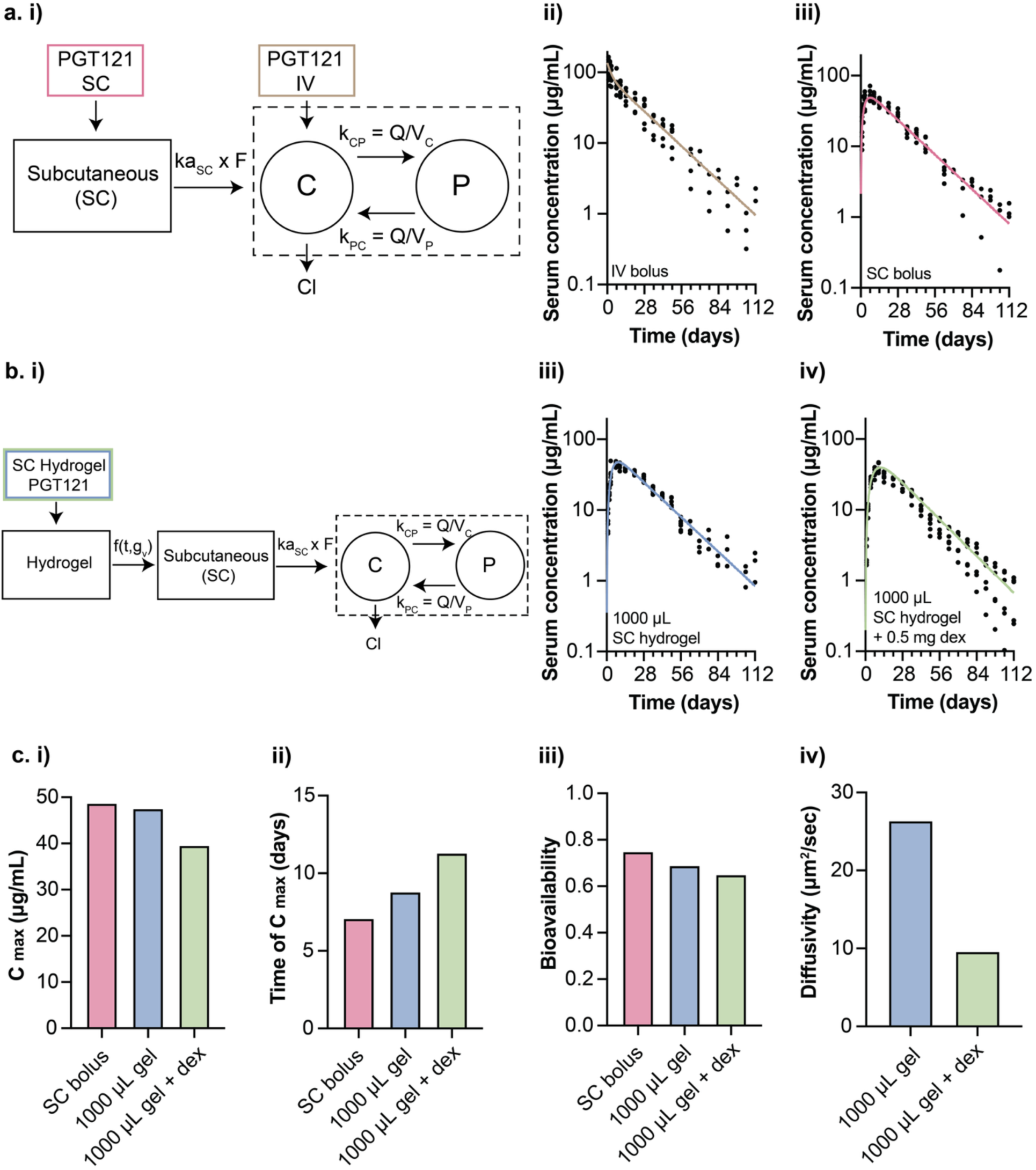
Multicompartmental modeling of mAb release from hydrogels in the subcutaneous space. **a. i)** General model diagram for PGT121 when administered intravenously (IV) or subcutaneously (SC). The rectangular region within the dashed lines depicts two-compartment kinetics of PGT121 flow between the central (C) and peripheral (P) compartments. Rate parameters are depicted above the arrows. Multicompartment model fits for **ii)** IV and **iii)** SC PK. **b. i)** General model diagram for PGT121 when administered via a subcutaneous hydrogel where f(t, g_V_) denotes a general flow process for the hydrogel to be investigated based on the given gel volume, g_V_. Multicompartment model fits for **ii)** 1 mL SC hydrogel and **iii)** 1 mL SC hydrogel with 0.5 mg dex. Corresponding **c. i)** Cmax, **ii)** time of Cmax, **iii)** bioavailability, and **iv)** gel diffusion constant.

**Table 1.**
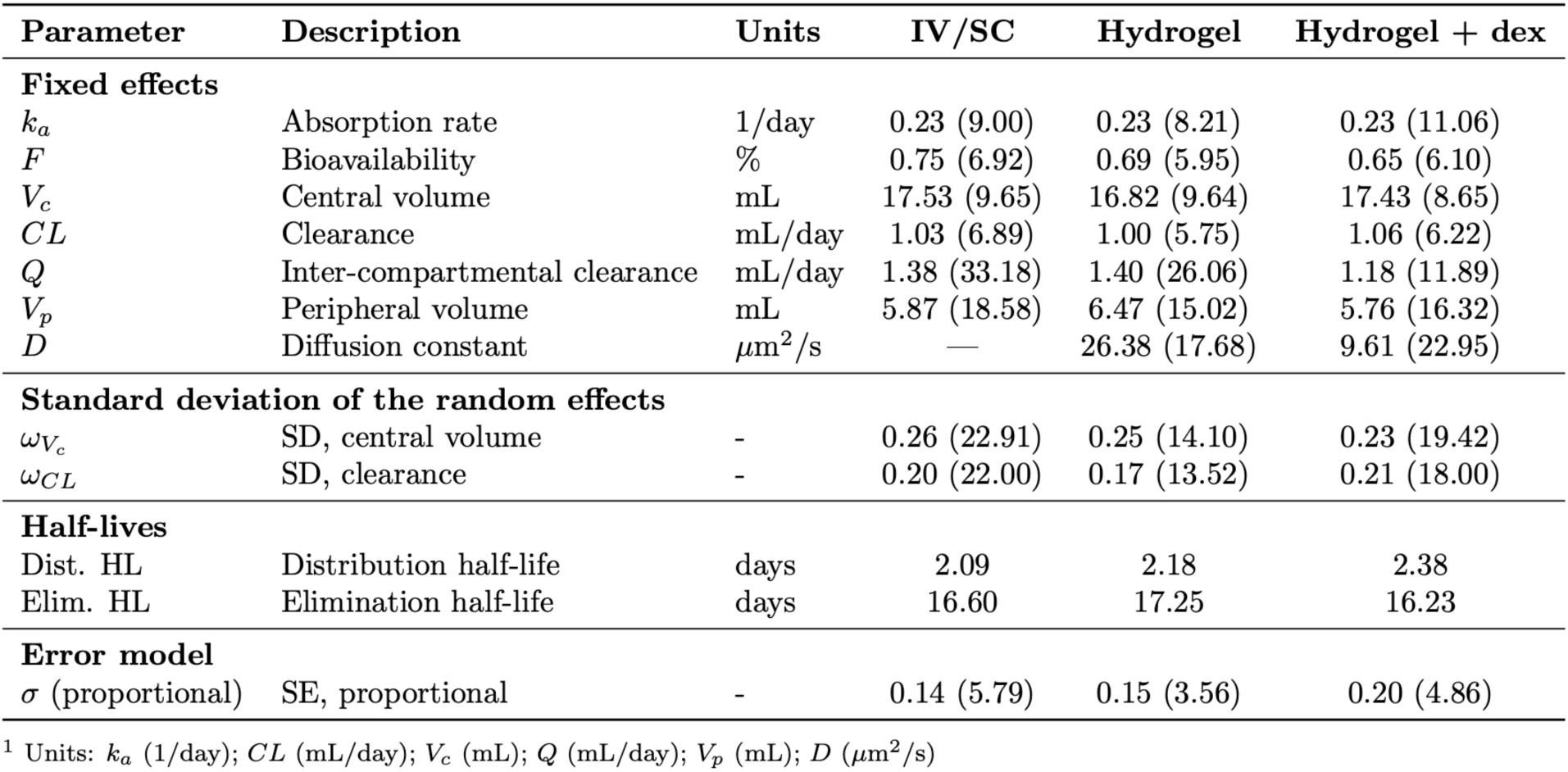
Population PK estimates (% Relative Standard Error) for PGT121. See Supplementary Discussion 2 for model descriptions. Symbol - denotes that the parameter was not estimated by the model. Standard errors for the PK model were estimated via linearization.

| Parameter | Description | Units | IV/SC | Hydrogel | Hydrogel + dex |
| --- | --- | --- | --- | --- | --- |
| **Fixed effects** |  |  |  |  |  |
| $k_a$ | Absorption rate | 1/day | 0.23 (9.00) | 0.23 (8.21) | 0.23 (11.06) |
| $F$ | Bioavailability | % | 0.75 (6.92) | 0.69 (5.95) | 0.65 (6.10) |
| $V_c$ | Central volume | mL | 17.53 (9.65) | 16.82 (9.64) | 17.43 (8.65) |
| $CL$ | Clearance | mL/day | 1.03 (6.89) | 1.00 (5.75) | 1.06 (6.22) |
| $Q$ | Inter-compartmental clearance | mL/day | 1.38 (33.18) | 1.40 (26.06) | 1.18 (11.89) |
| $V_p$ | Peripheral volume | mL | 5.87 (18.58) | 6.47 (15.02) | 5.76 (16.32) |
| $D$ | Diffusion constant | $\mu\text{m}^2/\text{s}$ | — | 26.38 (17.68) | 9.61 (22.95) |
| **Standard deviation of the random effects** |  |  |  |  |  |
| $\omega_{V_c}$ | SD, central volume | - | 0.26 (22.91) | 0.25 (14.10) | 0.23 (19.42) |
| $\omega_{CL}$ | SD, clearance | - | 0.20 (22.00) | 0.17 (13.52) | 0.21 (18.00) |
| **Half-lives** |  |  |  |  |  |
| Dist. HL | Distribution half-life | days | 2.09 | 2.18 | 2.38 |
| Elim. HL | Elimination half-life | days | 16.60 | 17.25 | 16.23 |
| **Error model** |  |  |  |  |  |
| $\sigma$ (proportional) | SE, proportional | - | 0.14 (5.79) | 0.15 (3.56) | 0.20 (4.86) |

<sup>1</sup> Units: $k_a$ (1/day); $CL$ (mL/day); $V_c$ (mL); $Q$ (mL/day); $V_p$ (mL); $D$ ( $\mu\text{m}^2/\text{s}$ )

Building on our modeling work in rats, we used the hydrogel PK model (**Fig. 9b i**) to simulate serum concentration profiles of HIV-neutralizing biologics in humans. Models assumed 2 mL subcutaneous injections, a subcutaneous bioavailability of 75%, and a 700 mg dose (corresponding to 10 mg/kg for a 70 kg individual). The terminal half-life of antibodies in humans was estimated using a combination of previously reported clinical PK data and scaling relationships of elimination half-lives in rodent models (**see Methods, Table 3**). Serum concentration profiles were simulated for PGT12, eCD4-Ig, and proteins that exhibit both longer and shorter elimination half-lives (**Fig. 10a-d**). PGT121.414.LS is a modified form of PGT121 with an extended half-life. IgM is a different antibody class with strong neutralization ability but a short in-vivo half-life. Four delivery strategies were modeled: IV injection, SC injection, a 2 mL hydrogel with dexamethasone, and a 2 mL hydrogel with an ideal diffusion coefficient of 2.6 µm²/s, based on FRAP analysis.

**Table 2.** Antibody panel for hydrogel staining.

| Stain | Dilution (1:X) | Dye | Source | Product Number |
| --- | --- | --- | --- | --- |
| anti-I-A/I-E | 800 | FITC | BioLegend | 107605 |
| anti-CD45 | 800 | AF700 | BioLegend | 103127 |
| anti-Ly6C | 400 | BV570 | BioLegend | 128029 |
| anti-XCR1 | 200 | AF647 | BioLegend | 148213 |
| anti-F4/80 | 200 | BV421 | BioLegend | 123137 |
| anti-Ly6G | 200 | BV711 | BioLegend | 127643 |
| anti-CD3e | 200 | PerCP-eFluor710 | Thermo Scientific | 46-0033-82 |
| anti-CD19 | 200 | PE-Cy7 | BioLegend | 115519 |
| anti-CD11c | 200 | PE | BioLegend | 117307 |
| anti-CD11b | 100 | BV510 | Fisher Scientific | 50-112-9846 |
| anti-NK1.1 | 100 | BV605 | BioLegend | 108753 |

**Table 3.** Input PK parameters for human-scale-up modeling.

| **mAb** | **Cl** | **Vc** | **Q** | **Vp** | **F** | **ka** | **Elimination Half-life (days)** |
| --- | --- | --- | --- | --- | --- | --- | --- |
| **PGT121** | 0.305 | 3.998 | 0.735 | 5.039 | 0.75 | 0.384 | 23 |
| **Theoretical eCD4-Ig** | 0.600 | 3.998 | 0.735 | 5.039 | 0.75 | 0.384 | 14 |
| **Theoretical IgM** | 0.543 | 3.998 |  |  | 0.75 | 0.384 | 5 |
| **PGT121.414.LS** | 0.062 | 2.942 | 0.541 | 3.023 | 0.75 | 0.277 | 69 |
| Units: ka (1/day); CL (L/day); Vc (L); Q (L/day); Vp (L) |  |  |  |  |  |  |  |

While the PK benefits of hydrogel administration are less evident for extended half-life bnAbs such as PGT121.414.LS (t_½ elim_ = 69 days) (**Fig. 10a, SI Fig 11**), for parental-type PGT121 (t_½ elim_ = 23 days) SC dosing via a hydrogel reduces C_max_ and allows for mimicking IV dosing in the terminal phase (**Fig. 10b**). When administered via an ideal PNP hydrogel material (D = 2.6 µm²/s) we see a further reduction in C_max_ and a 1.2-fold increase in time above an arbitrary therapeutic threshold (10 µg/mL) as compared to IV dosing (**Fig. 10e**). The advantages of hydrogel-based delivery are especially pronounced for shorter half-life antibodies such as eCD4-Ig (t_½ elim_ = 13.6 days) and IgM (t_½ elim_ = 5.1 days). For eCD4-Ig, hydrogels with diffusion coefficients of 9.6 µm²/s and 2.6 µm²/s extended time above the therapeutic threshold by 1.12- and 1.28-fold, respectively, compared to IV dosing (**Fig. 10e**). The effect was even more substantial for IgM, with corresponding fold increases of 1.38 and 1.68, highlighting the potential of PNP hydrogels to significantly enhance exposure for rapidly cleared biologics (**Fig. 10e**). Overall, these simulations highlight the potential of hydrogel depot technologies to enhance the PK of biologics in humans, particularly for short-lived antibodies or fusion proteins that would otherwise require frequent dosing.

## 3. Conclusion

In this study, we investigated the use of a subcutaneous PNP hydrogel depot technology for the extended delivery of HIV bnAbs. PNP hydrogels are a promising platform for controlled biologic release as they slow cargo diffusion, are easily injected, and form robust depots in the subcutaneous space. In-vivo PK studies in mice and rats demonstrated that PNP hydrogels deliver functional antibody over prolonged time-frames; however, the observed benefit of a hydrogel depot on the overall PK is dependent on both hydrogel volume and bnAb elimination half-life. Notably, in-vivo release was faster than predicted from in-vitro characterization. Flow cytometry analysis revealed that co-delivery of bnAbs with an anti-inflammatory agent reduces active cargo transport out of the depot via cellular uptake, thereby extending bnAb release. Hydrogel diffusivity values from rat PK data were incorporated into PK models to predict human serum profiles of clinically relevant HIV bnAbs. These models suggest that hydrogel depots may significantly improve the PK of biologics in humans, especially for biologics with short half-lives that would otherwise require frequent dosing. Collectively, these findings support the development of next-generation hydrogel depot systems for stabilizing and gradually releasing bnAbs, advancing strategies for long-term pre-exposure immunoprophylaxis against infectious diseases.

## 4. Materials and Methods

### 4.1 Materials

Unless otherwise specified, all reagent-grade chemicals and solvents were purchased from Sigma-Aldrich or Fisher Scientific and used without further purification. Human IgG (Cat. No. 340-21, Lot No. 07J4627) was obtained as a lyophilized powder from Medix Biochemica. HyPure™ Cell Culture Grade Water was sourced from Cytiva, and Phosphate Buffered Saline (Cat. No. 10010-023) was obtained from Gibco. The anti-HIV antibody-like protein eCD4-Ig (1v234-Emm-009) was generously provided by the Farzan Lab, and PGT121 was received from Just Biotherapeutics.

### 4.2 PNP Gel Synthesis

Polymer nanoparticle (PNP) hydrogels are formed from the mixing of PEG-PLA nanoparticles and hydrophobically modified (hydroxypropyl)methylcellulose.

#### 4.2.1 Poly(ethylene)-block-poly(lactic acid) Synthesis

Poly(ethylene glycol)-block-poly(lactic acid) (PEG-PLA) block copolymers were synthesized by ring opening polymerization as described in the methods of previously published work (*55*).

Polymer molecular weight and dispersity were characterized by SEC using DMF as an eluent (22.5–27.5 kDa (5 kDa PEG, 17.5–22.5 kDa PLA) with Đ < 1.2).

#### 4.2.2 PEG-PLA Nanoparticle Synthesis

PEG-PLA core-shell nanoparticles were formed via nanoprecipitation as described in previously published work (*55*). In short, PEG–PLA (50 mg) was dissolved in a mixture of acetonitrile and DMSO at a 75:25 ratio (1 mL total volume). This polymer solution was then slowly added dropwise into 10 mL of ultrapure water under vigorous stirring (600 rpm). The resulting nanoparticle suspension was concentrated by centrifugation using an Amicon Ultra-15 filter with a 10 kDa molecular weight cutoff. The nanoparticles were subsequently resuspended in PBS or the appropriate mAb buffer to a concentration of 20 wt.%. The hydrodynamic diameter of the nanoparticles, which ranged from 30 to 35 nm with a PDI of less than 0.2, was determined using dynamic light scattering (DLS) with a DynaPro II plate reader (Wyatt Technology).

#### 4.2.3 Dodecyl-Modified (hydroxypropyl)methylcellulose (HPMC-C12) Synthesis

Dodecyl-modified (hydroxypropyl)methyl cellulose (HPMC-C12) was prepared following methods described in previously published work (*55*). The resulting HPMC-C12 product was freeze-dried and then dissolved in phosphate-buffered saline (PBS) or the appropriate mAb buffer to create a 6 wt.% stock solution. Proton nuclear magnetic resonance (H-NMR) analysis was performed on the starting materials—Hypromellose (USP grade, 1 g) and dodecyl isocyanate (99%, 125 µL, 0.52 mmol)—as well as on the synthesized HPMC-C12, as documented in prior studies (*46, 51*). H-NMR results showed that the degree of dodecyl modification was 8.5 wt.%, which is close to the theoretical maximum of 10 wt.% for HPMC-C12 (*46*).

#### 4.2.4 Polymer Nanoparticle Hydrogel Synthesis

Supramolecular PNP hydrogels were prepared following methods described in previously published work (*55, 68*). In short, HPMC-C12 stock solutions and PEG-PLA nanoparticle stock solutions were mixed with PBS or the appropriate mAb buffer at specified concentrations using elbow mixing. The notation for PNP hydrogel formulations is defined as HPMC-C12 wt.% – NP wt.%. A PNP-2-10 hydrogel contains 2 wt.% HPMC-C12 and 10 wt.% nanoparticles, making up a total of 12 wt.% solids, with the remaining portion consisting of mAb cargo or buffer. To prepare a PNP-2-10 hydrogel, 334 mg of HPMC-C12 stock solution was placed in a 3 mL luer lock syringe, while 500 µL of nanoparticle stock solution was combined with 166 µL of mAb or buffer and loaded into another 3 mL luer lock syringe. The two syringes were connected using an elbow fitting and mixed for more than 100 cycles until a uniform hydrogel was formed.

Dexamethasone was added to the hydrogel by first preparing a stock solution of 100 mg/mL dex in DMSO. Dexamethasone in DMSO was first combined with buffer, then mAb, and finally nanoparticle stock solution. The combined mixture was loaded into a 3 mL luer lock syringe which was then mixed with HPMC-C12 as previously described.

### 4.3 PNP Gel Characterization

#### 4.3.1 Rheological Characterization

Rheological measurements were conducted at 4 °C, 25 °C, and 37 °C using a stress-controlled TA Instruments DHR-2 rheometer equipped with a serrated parallel plate of 20 mm diameter and a gap of 500 µm. Frequency sweeps were carried out at a strain of 1%, ensuring they remained within the linear viscoelastic range.

#### 4.3.2 Fluorescent Recovery after Photobleaching

PNP hydrogels were prepared with the soluble fraction containing FITC-labeled human IgG at a concentration of 5 mg/mL. 100 uLs of hydrogel was deposited onto a glass slide and imaged using a confocal LSM780 microscope. Fluorescence recovery after photobleaching (FRAP) was performed by bleaching a circular region of interest with a high-intensity 488 nm laser. Following photobleaching, fluorescence recovery was monitored for 200 seconds. The normalized recovery data were analyzed using previously reported fitting methods (*69*) to extract diffusivity values. Diffusivities of fluorescent probes in PBS were estimated using the Stokes–Einstein equation, assuming a hydrodynamic radius of 5.2 nm for hIgG and viscosity for PBS of 0.887 cP.

#### 4.3.3 In-vitro IgG Release

PNP hydrogels containing AF647-labeled human IgG in the soluble phase (5 mg/mL) were prepared for release studies. Borosilicate glass capillary tubes (2.7 mm inner diameter) were cut into 4-inch segments, and one end of each tube was sealed using epoxy resin. A volume of 100 µL of the hydrogel was dispensed into the sealed end of each tube. To eliminate trapped air, the tubes were centrifuged at 1000 rpm. Subsequently, 400 µL of phosphate-buffered saline (PBS) was added atop the gel in each tube. At designated intervals, the PBS was completely removed and replaced with fresh buffer. The collected PBS samples were analyzed for IgG content by fluorescence with excitation at 640 nm and emission at 670 nm. Quantification was performed using a calibration curve generated from known concentrations of AF647-labeled human IgG in PBS. During incubation between sampling timepoints, the tubes were stored at 37°C and sealed with parafilm to minimize evaporation.

### 4.4 Animal Studies

Animal studies were performed with the approval of the Stanford Administrative Panel on Laboratory Animal Care (APLAC-32109) and were in accordance with National Institutes of Health guidelines.

#### 4.4.1 eCD4 Ig Delivery in Mice

Female B6.Cg-Fcgrttm1Dcr Prkdcscid Tg(FCGRT)32Dcr/DcrJ mice (Jackson Laboratory, Strain No. 018441), aged 12–14 weeks, were administered anti-HIV antibody-like protein eCD4-Ig (1v234-Emm-009) either intravenously (via retro-orbital injection) or subcutaneously under brief isoflurane anesthesia. All antibody formulations were prepared at a dose of 360 µg eCD4-Ig per mouse. Mice in the gel group (n=4) received 200 µL of gel containing eCD4-Ig at a concentration of 1.8 mg mL⁻¹, while bolus mice were given 150 µL of eCD4-Ig at 2.4 mg mL⁻¹ in PBS via either retro-orbital (n=6) or subcutaneous (n=6) injection. Hydrogels were prepared using a 6 wt.% stock solution of HPMC-C12 in PBS, a 20 wt.% nanoparticle stock solution, and the appropriate concentration of eCD4-Ig in PBS. The samples were mixed by loading HPMC-C12 and eCD4-Ig + NP stock solutions into separate syringes, which were then connected using a luer lock elbow fitting to facilitate mixing through repeated passage. This method allowed for gentle encapsulation of the antibody and resulted in pre-loaded syringes ready for injection. Serum samples were collected from the tail vein at specified intervals to assess eCD4-Ig concentrations by ELISA. Samples used for analysis were collected at 1, 2.5, 4, 7, 10, and 14 days.

#### 4.4.2 PGT121 Delivery in Mice

Female B6.Cg-Fcgrttm1Dcr Prkdcscid Tg(FCGRT)32Dcr/DcrJ mice (Jackson Laboratory, Strain No. 018441), aged 12–14 weeks, were administered anti-HIV antibody PGT121 (Just Biotherapeutics) either intravenously (via retro-orbital injection) or subcutaneously under brief isoflurane anesthesia. All antibody formulations were prepared at a dose of 1 mg PGT121 per mouse. Mice in the gel group (n=7) received 200 µL of gel containing PGT121 at a concentration of 5 mg mL⁻¹, while bolus mice were given 150 µL of PGT121 at 6.66 mg mL⁻¹ in acetate buffer via either retro-orbital (n=11) or subcutaneous (n=8) injection. Hydrogels were prepared as previously described. Serum samples were collected from the tail vein at specified intervals to assess PGT121 concentrations by ELISA. Samples used for analysis were collected at 1, 4, 7, 10, 14, and 21 days.

#### 4.4.3 PGT121 Delivery in Rats

Male Sprague Dawley-Rag2em2heraIl2rgem1hera/HblCrl (SRG) rats (Charles River, Strain No. 707), age 12 weeks, were administered anti-HIV antibody PGT121 (Just Biotherapeutics) either intravenously (via tail vein injection) or subcutaneously (with or without hydrogel) under brief isoflurane anesthesia. All antibody formulations were prepared at a dose of 2.3 mg PGT121 per rat. The experiment consisted of 5 groups (all n = 6): one receiving PGT121 via IV, one receiving PGT121 via SC, and the 3 remaining groups receiving injections of hydrogels of volumes of 250, 500, and 1000 µL. Hydrogels were prepared as previously described. Serum samples were collected via tail vein. All animals were sampled at days 0 (pre-bleed), 1, 2, 3, 7, 10, 14, and then weekly through day 112. Rats were weighed weekly.

To assess the impact of immunosuppressant cargo on PK, a later experiment added two additional hydrogel groups (both n=6) where rats received 1000 µL hydrogel injections containing 2.3 mg PGT121 and either 0.5 mg or 1 mg of dexamethasone (dex) (**SI Fig 12**). Animals were age matched, and serum samples were collected via tail vein at days 0 (pre-bleed), 1, 2, 3, 7, 10, 14, and then weekly through day 112. Rats were weighed weekly.

#### 4.4.4 hIgG Delivery in Mice

Female C57BL/6NCrl mice (Charles River, Strain No. 027), age 12 weeks, were administered polyclonal human immunoglobulin G (hIgG) from Medix Biochemica (Cat No: 340-21, Lot: 07J4627) either as a subcutaneous bolus or via a subcutaneously injected hydrogel. All animals (all n=5) received 1 mg of hIgG. Hydrogels were prepared with hIgG alone as well as hIgG co-delivered with either dexamethasone (dex) at a low (0.02 mg/mouse) or high dose (0.2 mg/mouse) or Granulocyte-Macrophage Colony-Stimulating Factor (GM-CSF) at a dose of 3 µg/mouse. Hydrogels containing hIgG as well as immunostimulatory cargo were prepared using standard methods previously described. Serum samples were collected from the tail vein at specified intervals to assess hIgG concentrations by ELISA. Samples used for analysis were collected at 1, 2, 4, 7, 10, and 14 days.

#### 4.4.5 Quantification of Immune Cell Infiltration via Flow Cytometry

Female C57BL/6 mice (Charles River, Strain No. 027), age 12 weeks, were given subcutaneous injections of hydrogel containing either 1 mg of hIgG, 1 mg of hIgG + 0.2 mg dex, or 1 mg of hIgG + 3 µg GMCSF. Two days after injection, mice were euthanized by carbon dioxide asphyxiation. The hydrogel depots were then excised from the subcutaneous space and placed into microcentrifuge tubes containing 750 μL of FACS buffer (PBS 1X, 3% heat-inactivated FBS, 1 mM EDTA). To create single-cell suspensions, the hydrogels were mechanically disrupted using Kimble BioMasherIIs (DWK Life Sciences) tissue grinders. The resulting suspensions were filtered through 70 μm cell strainers (Celltreat, 229484) into 15 mL Falcon tubes, centrifuged at 400 RCF for 4 minutes, and resuspended in 300 μL of 1X PBS. Cells were counted using live-dead acridine orange/propidium iodide stain (Vitascientific, LGBD10012) on a Luna-FL dual fluorescence cell counter (Logos Biosystems).

Following counting, one million live cells per sample were transferred to a 96-well conical bottom plate (Thermo Fisher Scientific, 249570). Initially, cells were stained with 100 μL of Live/Dead Fixable Near-IR (Thermo Fisher Scientific, L34975) for 30 minutes at room temperature. The staining was stopped by adding 100 μL of FACS buffer, followed by centrifugation at 935 RCF for 2 minutes. Next, cells were incubated on ice for 5 minutes with 50 μL of anti-mouse CD16/CD32 (1:50 dilution; BD Biosciences, 553142) before applying 50 μL of a full antibody stain for 30 minutes on ice. After staining, cells were washed with 100 μL of FACS buffer, centrifuged at 935 RCF for 2 minutes, and resuspended in 60 μL of FACS buffer.

The samples were analyzed using the BD FACSymphony A5 SORP at the Stanford Shared FACS Facility, and data were processed in FlowJo using gating strategies available in the SI (**SI Fig 8**). The antibody panel for hydrogel staining included the following:

### 4.5 Pharmacokinetic (PK) Analysis

Concentration of protein in serum was quantified via binding ELISA. Ability to neutralize virus was quantified via neutralization assays.

#### 4.5.1 eCD4-Ig ELISA

ELISAs were used to quantify eCD4-Ig concentration in serum. 25 µL per well of 9e9 (3 µg/mL in PBS) was added to 96-well high-binding half-area plates (Corning™ 3690), and the plates were incubated overnight at 4°C. After incubation, the plates were washed twice with PBS-T (150 µL per well) and blocked with 125 µL of 2% BSA in PBS for one hour at 37°C. Serum samples were diluted in 2% BSA to create a range of dilutions starting from 1:50 to 1:200, followed by serial two-fold dilutions to ensure detection occurred within the linear range. For the standard curve, 270 µL of BSA was mixed with 30 µL of serum (10x dilution), and eCD4-Ig was added at the desired concentration. After blocking, 50 µL of standards and samples were added to the wells and incubated at 37°C for one hour. A secondary antibody mixture was prepared by adding 1.2 µL of secondary antibody (Jackson ImmunoResearch, AB_2337591) to 6 mL of 2% BSA to achieve a 1:5000 dilution. After the primary antibody incubation, the plates were washed five times with PBS-T and incubated with 50 µL of the secondary antibody mix for one hour at 37°C. Following secondary antibody incubation, the plates were washed ten times with PBS-T, and 50 µL of TMB solution (Abcam ab171523) was added to each well. The plate was incubated at room temperature for 3–10 minutes, or until color development was visible. Reactions were stopped by adding 50 µL of TMB Stop Solution (1 N HCl), causing a yellow color change. The absorbance was then read at 450 nm using a plate reader.

#### 4.5.2 PGT121 ELISA

Fluorescent carboxylated xMAP (Luminex Corp) were covalently coupled to neutravidin and then bound to goat anti-mouse IgG biotin, followed by binding to anti-PGT121 idiotype (ID) antibody (9e9) as previously described (*22, 70*). Serum samples were pre-diluted at 1:10 and then subsequently diluted at 1:100 (for final 1:1000 dilution) or higher for accurate quantification of PGT121 levels. Each sample was tested at 3 or more separate dilutions and some samples were tested in at least 2 separate assays to confirm observed concentration. PGT121 was titrated as a standard curve on every assay plate and used to determine the observed concentration in each sample. The 5PL EC50, area under the curve (AUC), and fluorescence intensity (FI) of the highest concentration were tracked in Levey Jennings to ensure consistency of control performance over time. The lower limit of quantitation of the assay was 0.1221 ng/ml. Every assay contained 5 spiked diluent controls to assess standard curve accuracy. Negative controls include Blank (uncoupled) beads (negative control bead), blank well (beads + detection antibody), CH58 mAb (HIV Env-specific).

#### 4.5.3 hIgG ELISA

The concentration of hIgG in serum was quantified using an IgG (Total) Human ELISA Kit (ThermoFisher Catalog #BMS2091). Serum samples were diluted in 2% BSA at a dilution ratio of 1:2500. Two-fold serial dilutions were then performed to ensure detection occurred within the linear range. A standard curve was included on every plate and used to determine the observed concentration in each sample.

#### 4.5.4 HIV-1 Neutralization

The TZM-bl neutralization assay was performed as previously described (*71*). Briefly, serum samples were heat-inactivated for 30 minutes at 56°C and tested using a primary 1:40 dilution and a 5-fold titration series against HIV-1 Env pseudovirus X2088_c9 which is highly sensitive to neutralization by both eCD4-Ig and PGT121. Purified eCD4-Ig and PGT121 IgG were tested in parallel at a starting concentration of 10 μg/mL with a 5-fold titration series. Estimated serum concentrations of eCD4-Ig and PGT121 were calculated by multiplying the determined ID50 titer of the respective serum sample and the determined IC50 concentration of the eCD4-Ig and PGT121 monoclonal antibody standard. Virus pseudotyped with the envelope protein of murine leukemia virus (MuLV) was used as negative control.

### 4.6 Population Pharmacokinetics (PK) Modeling

#### 4.6.1 One-compartment Modeling

A single-phase exponential decay compartmental model was fit to the IV bolus data to determine k_elim_. A one-compartment PK model was next fit the SC bolus data and SC gel data to determine k_abs_ utilizing k_elim_ fit from IV data. M_0_ was fixed based on initial dose, and F and V_d_ were allowed to vary. The differential equations and analytical solutions for the one-compartment model are provided in the supplementary material (**Supplementary Discussion 1**).

#### 4.6.2 Multicompartment Population PK Modeling

Mulitcompartment population PK models were developed to describe the serum PK of PGT121 in rats delivered subcutaneously via hydrogels. Initially, a two-compartment model with absorption, a standard model describing PK of monoclonal antibodies (*72*), was used to describe serum PK in mice receiving PGT121 via SC or IV, before being extended to include hydrogel cargo release (**Fig. 9a i, Fig. 9b i**). Serum concentrations were described using non-linear mixed effects models, where population-level (average) parameters were represented as fixed effects and inter-animal variation of these parameters were described by random effects. Population parameters were estimated by fitting the nonlinear mixed effects models using the method of maximum likelihood via the Stochastic Approximation Expectation Maximization (SAEM) algorithm, as implemented by Monolix version 2024R1 (*73*). Cargo release from hydrogel gel was modeled using approximations of Fickian diffusion equations for a sphere depending on the volume of the gel and a diffusion constant (D, µm²/s) (*53, 74*). In addition to the serum PK parameters, the D parameter was also fitted using the serum concentration data. Full details on model fitting, including differential questions are provided in the supplementary material (**Supplementary Discussion 2**).

#### 4.6.3 Clinical Compartment Modeling

Clinical serum PK simulations using hydrogel delivery were performed using the hydrogel population PK model described in Supplementary Discussion 2 parameterized with clinical settings. A dose of 700 mg was used based on a 10 mg/kg dosing for an average 70 kg person. Four administration routes were used: IV, SC, hydrogel+dex with a diffusion rate of 9.6 *μ*m^2^/s based on in-vivo rat PK, and hydrogel+dex with a diffusion rate of 2.62 *μ*m^2^/s based on in-vitro FRAP experiments.

Two-compartment PK parameters for PGT121 and the modified PGT121.414.LS were derived from previous PK assessment of these antibodies (*19, 75*). The two-compartment parameters for the theoretical eCD4-Ig were determined based on the wild-type PGT121 but with 2-fold shorter half-life as observed in the mice experiments. For the theoretical IgM PK, data in healthy participants remains limited (*76*), but a one-compartment model was used based on a clinical experiment of IgM where rapid equilibrium was observed (*77*). The half-life for IgM was observed to be 5.1 days and the serum volume was assumed to be equivalent to the PGT121 value for comparability. The SC absorption rate was set to be equivalent for PGT121, eCD4-Ig, and IgM based on the PGT121 value. We considered a scenario in which bioavailability was assumed to be 75%. Input PK parameters were the following:

## 5. Statistical methods

For in-vivo experiments, animals were cage blocked, and Mead’s Resource Equation was applied to determine the appropriate sample size to ensure that additional subjects would have minimal impact on statistical power. Comparisons between two groups were performed using a two-tailed Student’s t-test, while comparisons involving more than two groups were analyzed using a 1-way analysis of variance (ANOVA) with Tukey’s correction for multiple comparisons in GraphPad PRISM. Results were considered statistically significant if p < 0.05.

## 6. Acknowledgements

The authors are grateful for financial support from the Center for Human Systems Immunology with the Gates Foundation (OPP1113682; INV-010680), the Vaccine & Immunology Statistical Center with the Gates Foundation (INV-032929), the Gates Foundation (OPP1211043; INV-027411 and INV-036842), and the NIH (5R01AI154989-06). Flow cytometry was performed on an instrument in the Stanford Shared FACS Facility that utilizes NIH S10 Shared Instrument Grant (1S10OD026831-01).

C.M.K. was supported by a Stanford Graduate Fellowship and the Stanford Bio-X William and Lynda Steere Fellowship. C.K.J. and J.Y. acknowledge support from National Science Foundation (NSF) Graduate Research Fellowships. P.G., O.M.S., and C.M.W. were supported by both NSF Graduate Research Fellowships and Stanford Graduate Fellowships. L.T.N. received support from a Stanford Graduate Fellowship. N.E. was supported by an NSF Graduate Research Fellowship (DGE-2146755), and A.I.D. was supported by a Schmidt Science Fellows Award. E.L.M. received support from the NIH Biotechnology Training Program (T32-GM008412), and V.M.D. is supported by K01EB033870. S.C.W. was supported by the Sarafan ChEM-H Chemistry/Biology Interface Training Program and the NSF Graduate Research Fellowship.

## 7. Author Contributions

C.K.J., C.M.K, and E.A.A. conceived of the idea. C.K.J. led the experimental work, performed the bulk of data analysis, and authored the manuscript. C.M.K designed and conducted initial studies with eCD4-Ig and PGT121 in mice and performed mouse eCD4-Ig binding ELISAs. S.S wrote python scripts for one-compartment modeling. B.T.M and O.H. performed multi-compartmental modeling. P.G. performed FRAP experiments. E.L.M analyzed flow cytometry data. R.R. and M.P. performed neutralization experiments. N.L.Y and G.D.T performed PGT121 binding ELISAs. All other authors aided in experiments.

