## Supplemental Information for "Engineering Sustained-Release Broadly Neutralizing Antibody Formulations"

### Supplementary Figures

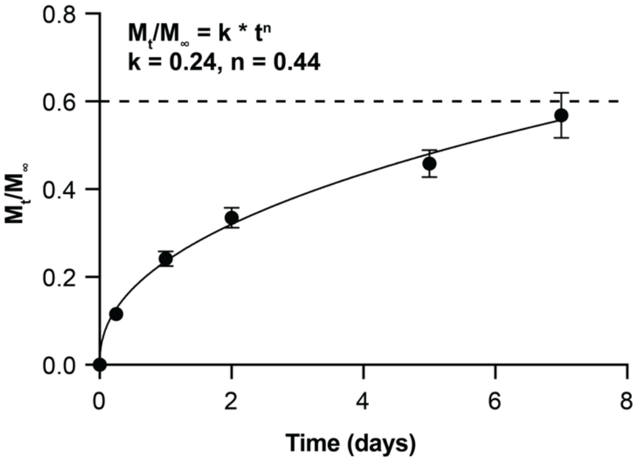

**SI Figure 1. Korsmeyer-Peppas fit of PNP in-vitro release data.** In-vitro release data from 2:10 PNP hydrogel (up to 60% release) is fit to Ritger-Peppas empirical equation where k is a constant and n is the diffusional exponent. A fit of n=0.44 suggests cargo release is dictated by Fickian diffusion.

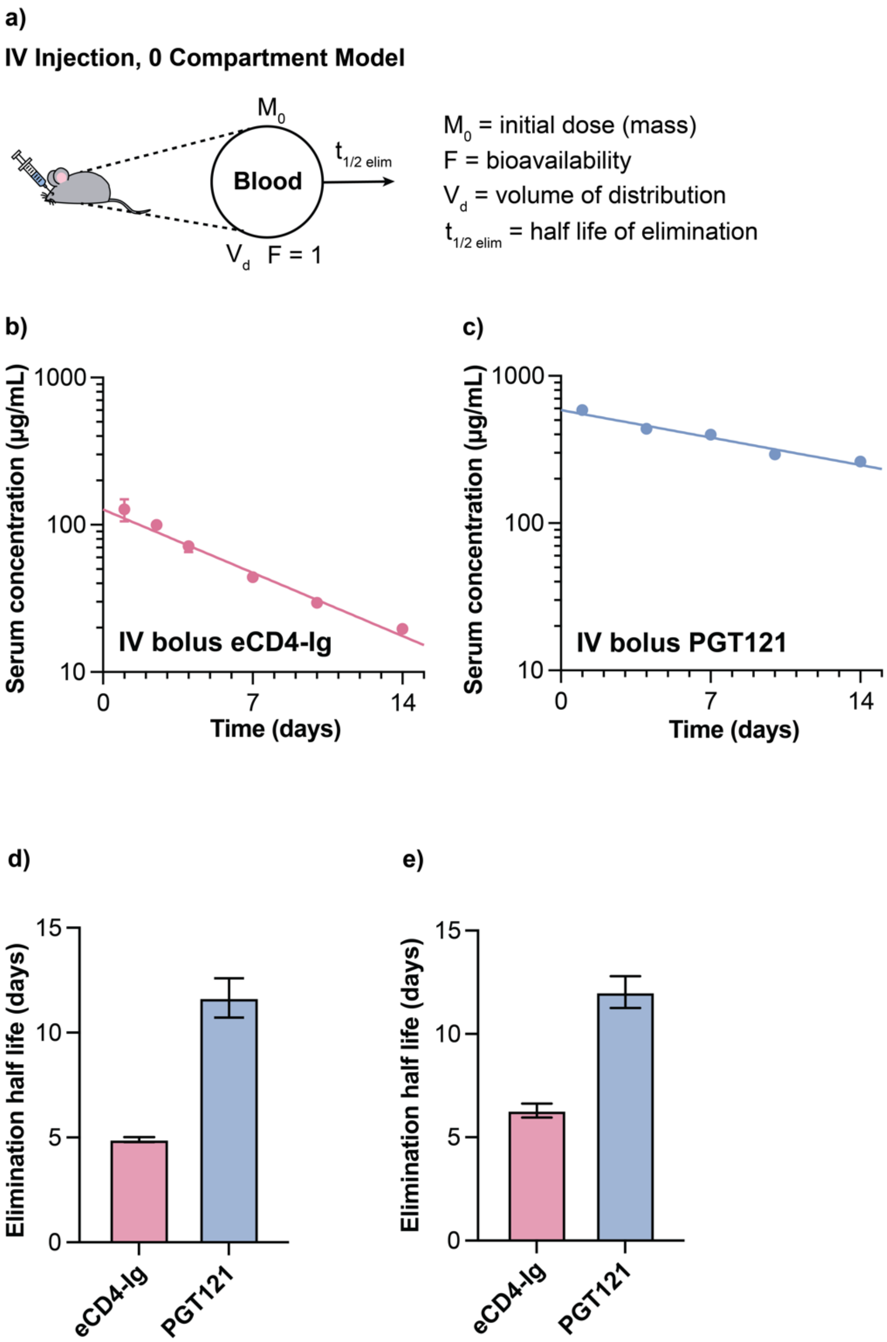

**SI Figure 2. Mouse eCD4-Ig and PGT121 IV bolus data fit to a single-phase exponential decay to obtain elimination half-life. a)** Schematic of zero-compartment model to fit IV data. **b)** eCD4-Ig IV bolus serum concentration profile (means with SEM) and **c)** PGT121 IV bolus serum concentration (means with SEM) profile with **d)** corresponding elimination half-life. Bar graph shows mean ± parameter fit uncertainty. **e)** Average elimination half-life for individual animal data fit to single-phase exponential decays. Bar graph shows mean ± SEM.

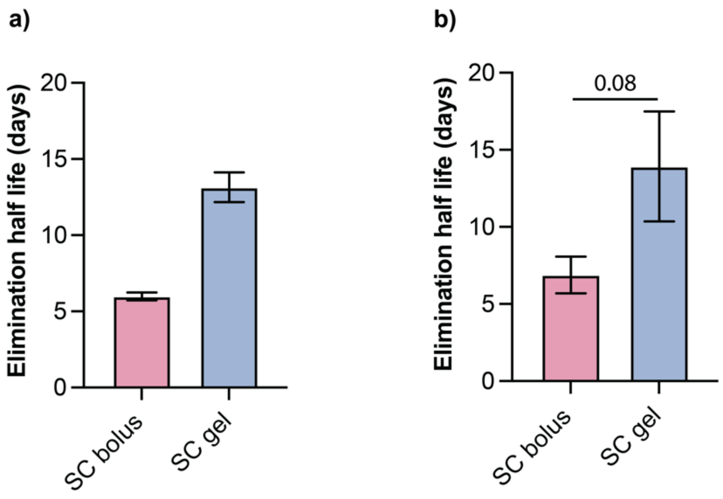

**SI Figure 3. Mouse eCD4-Ig SC bolus and SC gel data fit to a single-phase exponential decay to obtain elimination half-life. a)** Elimination half-life for serum concentration means and SEMs fit to a single-phase exponential decay. Bar graph shows mean ± parameter fit uncertainty. **d)** Average elimination half-life for individual animal data fit to single-phase exponential decays. Bar graph shows mean ± SEM. P values from student's t-test.

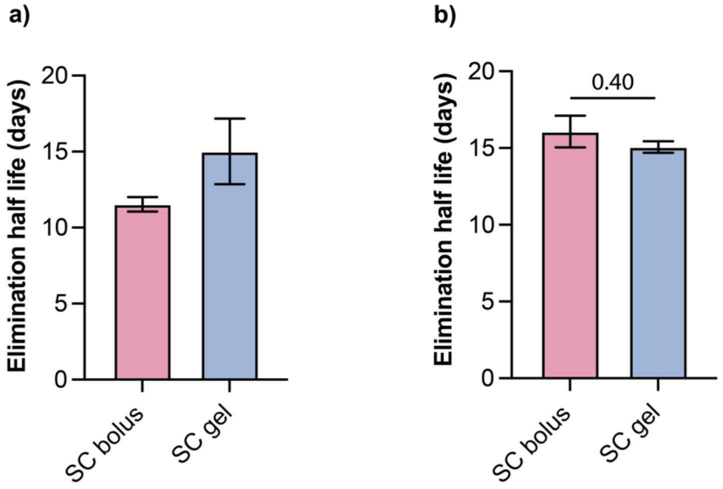

**SI Figure 4. Mouse PGT121 SC bolus and SC gel data fit to a single-phase exponential decay to obtain elimination half-life. a)** Elimination half-life for serum concentration means and SEMs fit to a single-phase exponential decay. Bar graph shows mean ± parameter fit uncertainty. **d)** Average elimination half-life for individual animal data fit to single-phase exponential decays. Bar graph shows mean ± SEM. P values from student's t-test.

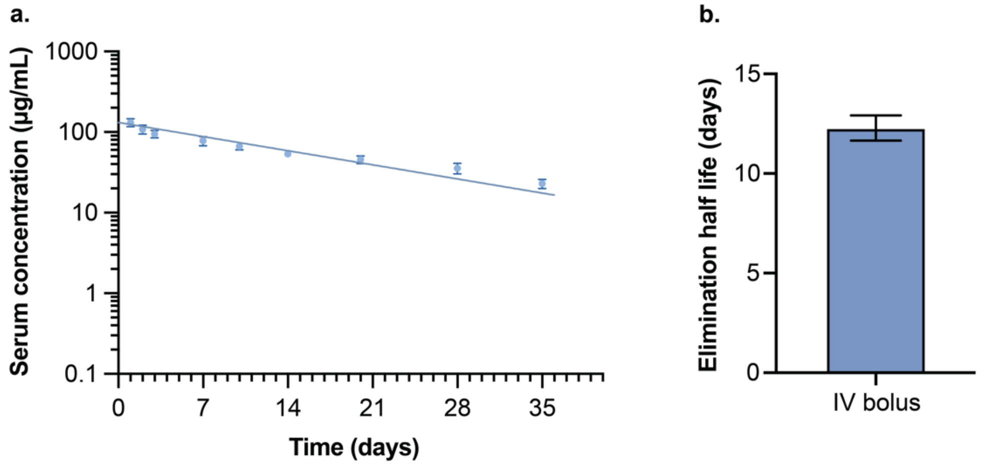

**SI Figure 5. Rat PGT121 IV bolus data fit to a single-phase exponential decay to obtain elimination half-life. a)** PGT121 IV bolus serum concentration profile (means with SEM) and **b)** average elimination half-life for individual animal data fit to single-phase exponential decays. Bar graph shows mean ± SEM.

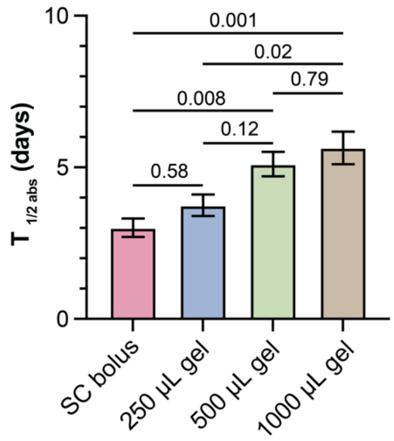

**SI Figure 6. Half-life of subcutaneous absorption ($T_{1/2abs}$) of SC gel and SC bolus delivery of PGT121.** Serum concentrations of PGT121 after SC bolus and SC gel administration were fit to obtain the half-life of subcutaneous absorption ($T_{1/2\ abs}$) using a one-compartment model. Bar graph shows mean ± SEM. P values from 1-way ANOVA with Tukey correction for multiple comparisons.

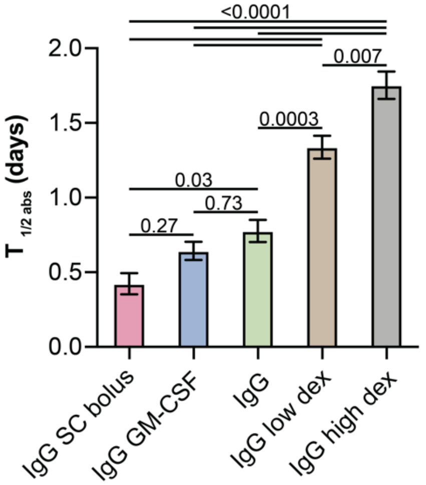

**SI Figure 7. Half-life of subcutaneous absorption ($T_{1/2abs}$) of SC gel, SC gel with dex, SC gel with GMCSF and SC bolus delivery of hIgG.** Serum concentrations of hIgG after SC bolus, SC gel, SC gel with dex, and SC gel with GMCSF administration were fit to obtain the half-life of subcutaneous absorption ($T_{1/2\ abs}$) using a one-compartment model. Bar graph shows mean ± SEM. P values from 1-way ANOVA with Tukey correction for multiple comparisons.

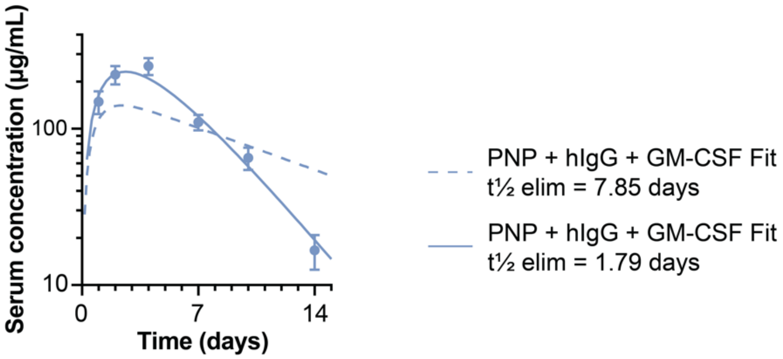

**SI Figure 8. One-compartment model fit for gel co-delivering hIgG and GM-CSF.** Serum concentrations of hIgG after SC administration of gels co-loaded with GM-CSF were fit to a one-compartment pharmacokinetic model. The model provided a poor fit when using the elimination half-life fit from SC bolus hIgG administration (7.85 days). Allowing the elimination half-life to vary improved the fit, yielding a best-fit elimination half-life of 1.79 days.

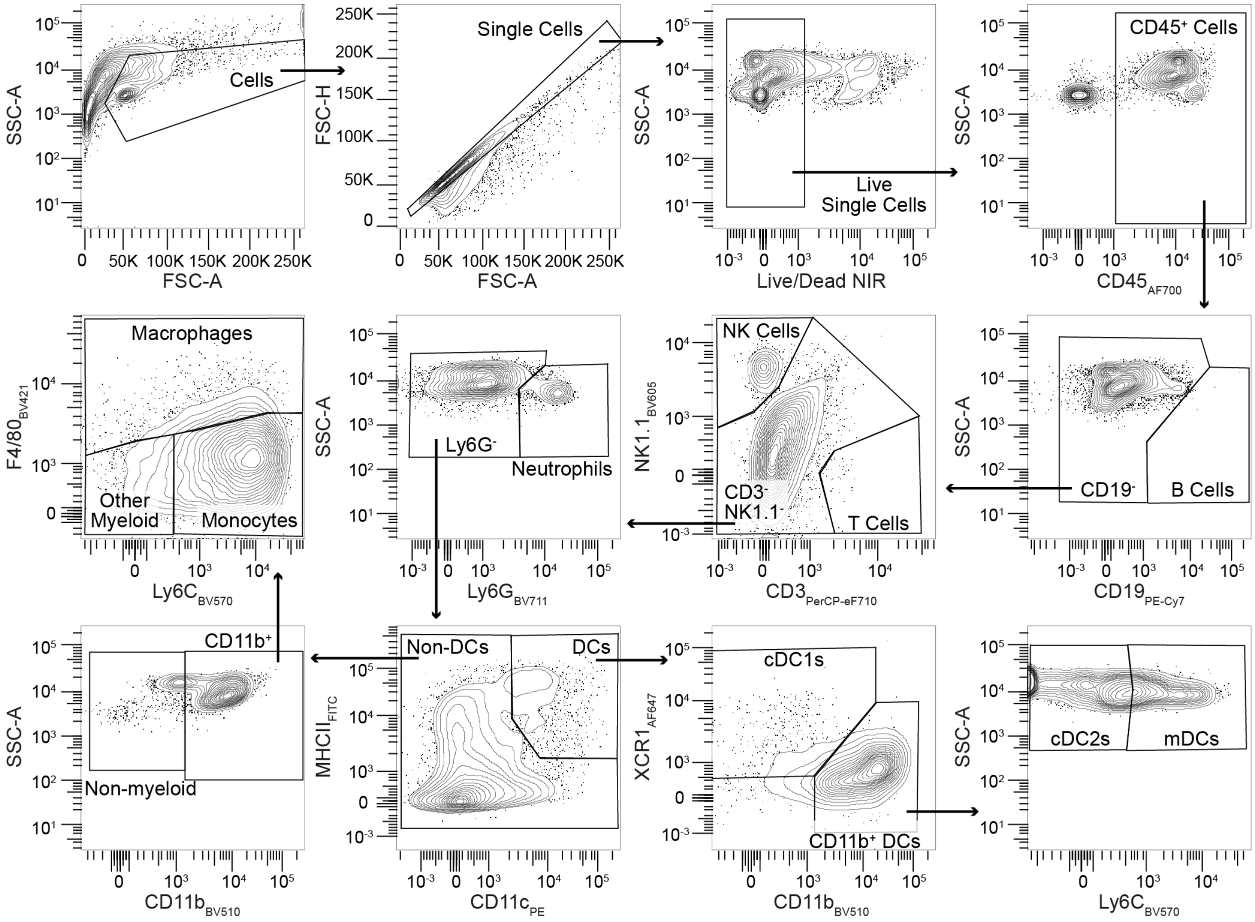

**SI Figure 9. Gating strategy for PNP hydrogels.** Shown with representative sample from PNP with IgG on day 2.

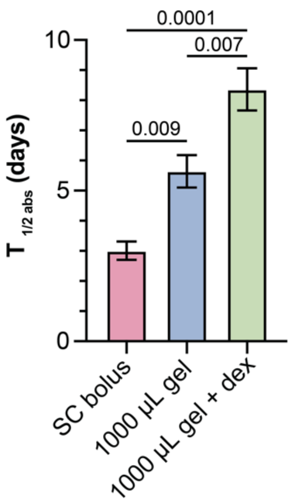

**SI Figure 10. Half-life of subcutaneous absorption ($T_{1/2abs}$) of SC gel, SC gel with dex, and SC bolus delivery of PGT121.** Serum concentrations of PGT121 after SC bolus, SC gel, and SC gel with dex administration were fit to obtain the half-life of subcutaneous absorption ($T_{1/2\ abs}$) using a one-compartment model. Bar graph shows mean ± SEM. P values from 1-way ANOVA with Tukey correction for multiple comparisons.

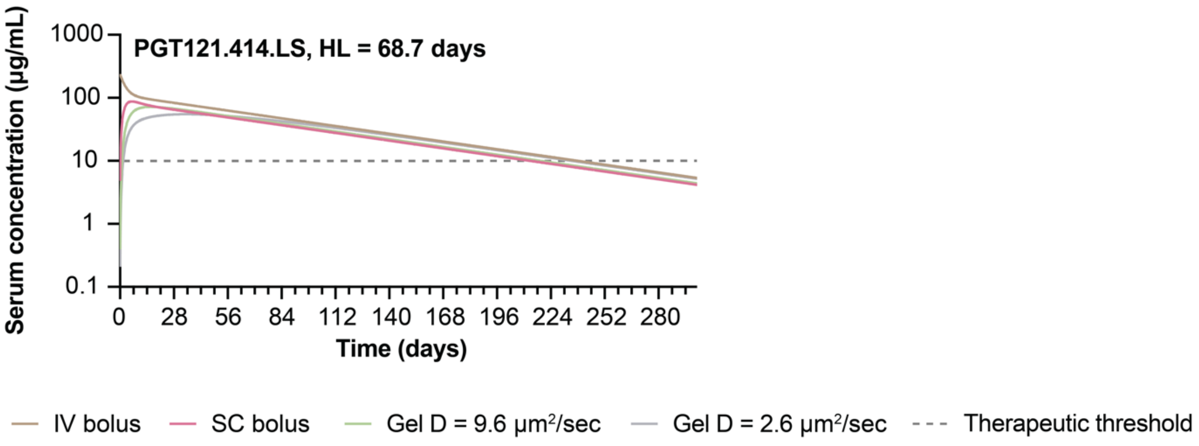

**SI Figure 11. Clinical PK simulations for delivery of PGT121.414.LS.** Modeled protein serum concentration for PGT121.4414.LS for IV injection, SC injection, injection of a 2 mL hydrogel with dexamethasone (D = 9.6 µm²/s), and injection of a 2 mL hydrogel with an ideal diffusion coefficient of 2.6 µm²/s. Therapeutic threshold is set at 10 µg/mL.

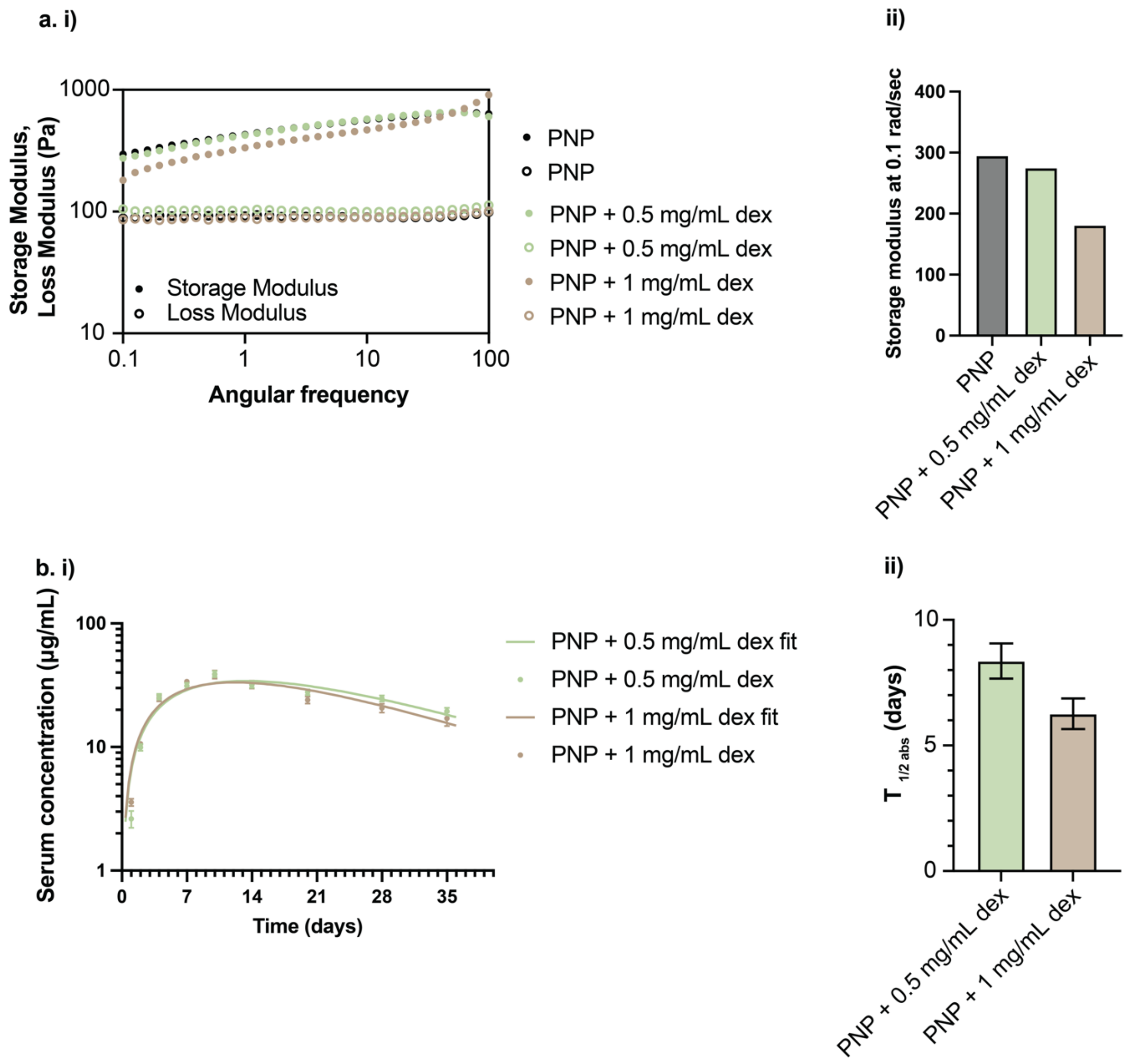

**SI Figure 12. Rheology and PK of high dexamethasone dose in rats. a. i)** Frequency sweeps of 2:10 PNP hydrogels that are empty and loaded with dex at 0.5 mg/mL and 1 mg/mL. Addition of higher dose dexamethasone (1 mg/mL) results in a weaker hydrogel network with a lower storage modulus. **ii)** Comparative storage modulus at 0.1 rad/sec. **b. i)** Serum concentrations of PGT121 after SC administration of low and high dose dex gels are fit to obtain **ii)** the half-life of subcutaneous absorption ($T_{1/2\ abs}$) using a one-compartment model. Higher dexamethasone doses result in faster release characterized by lower $T_{1/2\ abs}$. Bar graph shows mean ± SEM.

### Supplementary Discussion

#### 1. One-Compartment Pharmacokinetics modeling

The *in vivo* pharmacokinetics data were modeled using either a zero-compartment (IV bolus) or a one-compartment model (SC bolus / hydrogel groups). The kinetics of each process are assumed to be first order throughout.

For the IV bolus groups, the elimination half-life $\tau_e$ is obtained by fitting the bolus data to the following kinetic equation:

$$\frac{dM}{dt} = -k_{elim}M$$

where the elimination half-life $\tau_e$ and the elimination rate constant $k_{elim}$ satisfy the equation $\tau_e = \ln 2/k_{elim}$. The equation is solved with initial conditions $M(0) = M_0$, where $M_0$ is the total mass of cargo loaded into the hydrogels.

For the SC bolus and gel groups, there is a SC space compartment which may include a hydrogel depot to delay diffusive release. This compartment contains the initial cargo loading of mass $M_0$. Cargo is released from the SC space into the blood which has a nominal volume of distribution $V_d$. This process has a rate constant $k_{abs}$, which is related to the SC space absorption half-life as $\tau_{abs} = \ln 2/k_{abs}$. The cargo is eliminated from the blood at a rate of $k_{elim}$ where the elimination half-life is calculated as described earlier. The two-compartment model is mathematically expressed as:

$$\frac{dM_1}{dt} = -k_{abs}M_1$$

$$\frac{dM_2}{dt} = k_{abs}M_1 - k_{elim}M_2$$

where $M_1$ and $M_2$ are the mass of cargo in the SC space and blood respectively. The release and elimination rate constants and hence the corresponding half-lives are obtained as fit parameters from fitting the model to the PK data, assuming that the initial conditions are given by $M_1(0) = M_0$ and $M_2(0) = 0$.

Model fitting was carried out using a weighted least-squares residual optimization procedure using the *differential_evolution* fitting algorithm in Python. The residuals were weighted by the inverse of the standard errors on mean to incorporate uncertainty propagation.

#### 2. Multicompartment Population Pharmacokinetics Modeling

We developed a population PK model that described the PK of PGT121 in rats delivered subcutaneously via hydrogels. To that end, we initially constructed compartment models that describe PK of PGT121 in animals receiving the bnAb directly via either IV or SC administration.

We then extended the model to describe release of PGT121 from the hydrogels. The sections that follow provide details about the formulation of the candidate models and how these models were fitted to experimental data.

**Population model specification.** Serum PGT121 concentrations measured over time in multiple animals were described using nonlinear mixed effects models. For any given population PK model, population-level (average) parameters were represented as fixed effects and inter-animal variation of these parameters was described by random effects. Here, the standard deviations of the random effects were generically denoted by $\boldsymbol{\omega}$, with a subscript identifying the PK parameter to which the notation applies. A residual error model was specified to capture statistical properties of the outcome measure within each animal. PK models were fit to PGT121 serum concentrations assuming normally distributed error terms. We assumed that the residual error was proportional to the conditional expectation of the concentration, given the random effects. PK parameters were assumed to follow log-normal distributions with the exception of bioavailability, which was assumed to follow logit-normal distributions bounded between 0 and 100%. The variance-covariance matrix of the random effects was assumed to be diagonal with heterogeneous variances.

**Estimation of population parameters.** Population parameters, including the population mean and variance of each PK parameter and parameters of the error term variance, were estimated by fitting the nonlinear mixed effects models using the method of maximum likelihood, implemented using the Stochastic Approximation Expectation Maximization (SAEM) algorithm.

**Model fitting procedure**

Models were fit using the following three-step procedure that was designed to estimate fixed effects while gradually down-selecting random effects:

Step 1. Fit the model with the full random effects structure using a simulated annealing optimization algorithm.

Step 2. Set initial values to the results from Step 1 and re-fit the model using SAEM without simulated annealing.

Step 3. Assess and remove random effects based on whether the magnitude of the variance estimate approaches 0 (i.e., $< 1 \times 10^{-5}$) and/or percent relative standard error (%RSE) estimates > 50%.

In Step 1, simulated annealing allows the procedure to keep the explored parameter space large for longer during optimization of the log-likelihood function to escape local maxima and increase the chance of convergence towards a global maximum. Therefore, this approach was implemented to obtain good initial values for subsequent execution of the SAEM algorithm in Step 2 (*1*). It may also help retain parameters in the model that may otherwise be removed due to the optimization algorithm stopping at local maxima.

In Step 3 above, the %RSE for a given parameter was calculated as the standard error (times 100) divided by the estimated parameter value.

**PK model building**

Previous work on IgG monoclonal antibodies suggests that the PK for PGT121 may be properly described using a two-compartment model (*2, 3*). When applied in the context of monoclonal antibodies, a two-compartment model assumes that protein clearance is bi-phasic, allowing product distribution into two distinct compartments: the central (i.e., serum) and peripheral (i.e., tissue) compartments (**Fig. 9a i in the main text**). This model includes four PK parameters: the central volume ($\mathbf{V_c}$), clearance from the central compartment ($\mathbf{CL}$), peripheral volume ($\mathbf{V_p}$), and intercompartmental clearance ($\mathbf{Q}$).

The two-compartment PK model was first fit to observed serum concentrations from the IV and direct SC groups alone in order to verify whether it properly described serum concentrations of PGT121 over time. To include observations from the SC group, the two-compartment model was extended to include an absorption phase via a depot compartment to account for the gradual release of the antibody from the site of injection into the blood stream. To characterize the SC-route absorption, two parameters were used: the absorption rate, $\mathbf{k_a}$, and the bioavailability, $\mathbf{F}$.

Once a base PK model was established for PGT121 administered via IV and SC, the model served as the basis for the PK models that include the hydrogel route (see Figure 9 in main text). Modeling the hydrogel route with specific model structures and equations are described below.

**PK Model structure and equations**

The general two-compartment population PK model structure is depicted in Figure 9 in the main text. The model satisfied the following set of ordinary differential equations (ODEs):

$$\begin{aligned} G_t{}' &= -f(t, g_v), \\ D_t{}' &= f(t, g_v) - k_{a,sc}D_t \\ C_t{}' &= Fk_{a,sc}D_t - \left(k_e + k_{cp}\right)C_t + k_{pc}P_t, \\ P_t{}' &= k_{cp}C_t + k_{pc}P_t, \end{aligned}$$

where $\mathrm{G_t}$ denotes the mass of PGT121 in the hydrogel at time $\mathrm{t}$; $\mathrm{D_t}$ (depot) denotes the mass of PGT121 in the subcutaneous space at time $\mathrm{t}$; $\mathrm{C_t}$ (central) denotes the mass of PGT121 in the blood at time $\mathrm{t}$; and $\mathrm{P_t}$ (peripheral) denotes the mass of PGT121 in the tissues at time $\mathrm{t}$. The relationship between the rate parameters $\{\mathrm{k_e}, \mathrm{k_{cp}}, \mathrm{k_{pc}}\}$ and the PK parameters $\{\mathrm{V_c}, \mathrm{CL}, \mathrm{V_p}, \mathrm{Q}\}$ is as follows:

$$\begin{aligned} k_e &= \frac{CL}{V_c}, \\ k_{cp} &= \frac{Q}{V_c}, \\ k_{pc} &= \frac{Q}{V_p}. \end{aligned}$$

The distribution and elimination half-lives were computed, respectively, as

$$
\begin{aligned}
t_{\frac{1}{2},\alpha} &= \frac{log(2)}{\alpha}, \\
t_{\frac{1}{2},\beta} &= \frac{log(2)}{\beta},
\end{aligned}
$$

where $\alpha$ and $\beta$ describe the distribution and elimination phase bi-exponential rates, respectively, and were calculated as follows:

$$
\begin{aligned}
X &= k_{cp} + k_{pc} + k_e \\
Y &= k_{pc} \times k_e \\
\alpha &= \frac{-X + \sqrt{X^2 - 4Y}}{2} \\
\beta &= \frac{-X - \sqrt{X^2 - 4Y}}{2}.
\end{aligned}
$$

The function $\mathbf{f}(\mathbf{t}, \mathbf{g_v})$ describes the flow of PGT121 released by the hydrogels into the depot SC compartment. It depends on time $\mathbf{t}$ and on the hydrogel volume ($\mathbf{g_v}$) and its form is described below. In this analysis, we assumed that all PGT121 contained in the hydrogel was first released into the depot compartment (i.e, the subcutaneous space) and not directly into the blood stream via an alternative pathway. Under this assumption, the PGT121 absorption phase after release from the hydrogel is described by the same absorption rate and bioavailability parameters as the direct SC route.

Mathematical models of drug release from spheres have been described in detail using extensions of the Fickian diffusion equations (*4*). These models depend on two parameters: the sphere volume (via its radius, $\mathbf{r}$) and a diffusion coefficient ($\mathbf{D}$). An approximation of this mathematical system depends on the stage of release (i.e., how much drug remains) and is described accordingly using two equations (see equations (9)-(10) in Arifin, Lee, and Wang) (*5*):

$$
\frac{\mathbf{M_t}}{\mathbf{M_\infty}} = \begin{cases} 6\left(\frac{\mathbf{Dt}}{\boldsymbol{\pi}\mathbf{r^2}}\right)^{1/2} - \frac{\mathbf{3Dt}}{\mathbf{r^2}}, & \frac{\mathbf{M_t}}{\mathbf{M_\infty}} < \mathbf{c} \\ 1 - \frac{6}{\boldsymbol{\pi}^2}\exp\left(-\frac{\boldsymbol{\pi}^2\mathbf{Dt}}{\mathbf{r^2}}\right), & \frac{\mathbf{M_t}}{\mathbf{M_\infty}} > \mathbf{c}, \end{cases}
$$

where $\mathrm{M_t}/\mathrm{M_\infty}$ represents the cumulative fraction of PGT121 mass released by time $\mathrm{t}$. The constant $\mathrm{c}$ defines the upper/lower range of the intervals over which the proposed approximations are considered valid. It is suggested in Arifin, Lee, and Wang that c is around 0.4 for the early-stage approximation and 0.6 for the late-stage approximation.

To determine the value of c, we performed an investigation comparing the approximations for a range of gel volumes (250 - 1000 µL) and diffusion coefficients (0.5 - 50 µm/s) in the scope of this analysis and found that a smooth transition between the approximations occurs near a release fraction of 0.8 (**SI Fig 13**). We then solved for the time, $\mathrm{t_{0.8}}$, when the early approximation of $\mathrm{M_t}/\mathrm{M_\infty}$ reaches this threshold (i.e, $\mathrm{M_{t_{0.8}}}/\mathrm{M_\infty} = 0.8$), where

$$\mathbf{t_{0.8}} = \left(\frac{-6\sqrt{\frac{\mathbf{D}}{\boldsymbol{\pi}\mathbf{r^2}}} + \sqrt{36\frac{\mathbf{D}}{\boldsymbol{\pi}\mathbf{r^2}} - 12\frac{\mathbf{D}}{\mathbf{r^2}} \times \mathbf{0.8}}}{-6\frac{\mathbf{D}}{\mathbf{r^2}}}\right)^2.$$

Based on the initial mass in the gel, $G_0$, the mass in the gel at time $t$, $G_t$, can be computed from these fractional release models as

$$G_t = G_0\left(1 - \frac{M_t}{M_\infty}\right).$$

We incorporated this function into the PK models by calculating its derivative such that

$$f(t, g_v) = \begin{cases} 3G_0\left(\frac{D}{r^2} - \left(\frac{D}{\pi r^2 t}\right)^{1/2}\right), & t < t_{0.8} \\ -\frac{\pi^2 D}{r^2} G_t, & t > t_{0.8}, \end{cases}$$

where $r = \left(\frac{3g_v \times 10^9}{4\pi}\right)^{1/3}$ with $g_v$ given in μL and $r$ given in μm.

The two-compartment PK model with hydrogel route was then fit to all groups (either with or without dex). Initial conditions were taken from the base model fit to the IV and SC groups alone. The random effects were not re-evaluated when adding the hydrogel groups. While the radius of the gel was fixed depending on the gel volume, we fit the diffusion coefficient, $D$, given in $\mu m^2/s$, to these data assuming it was log-normally distributed and does not vary between animals (i.e., no random effect). In the model, the coefficient was converted to $\mu m^2$/day by multiplying by a factor 86,400 s/day. The dosing of PGT121 was 2.3 mg in all groups and the initial conditions were specified as follows in accordance with the administration route: for IV, $C_0 = 2.3$ mg; for SC, $D_0 = 2.3$ mg; and for hydrogel, $G_0 = 2.3$ mg. Sera concentrations were calculated as $C_t/V_c$ for model fitting.

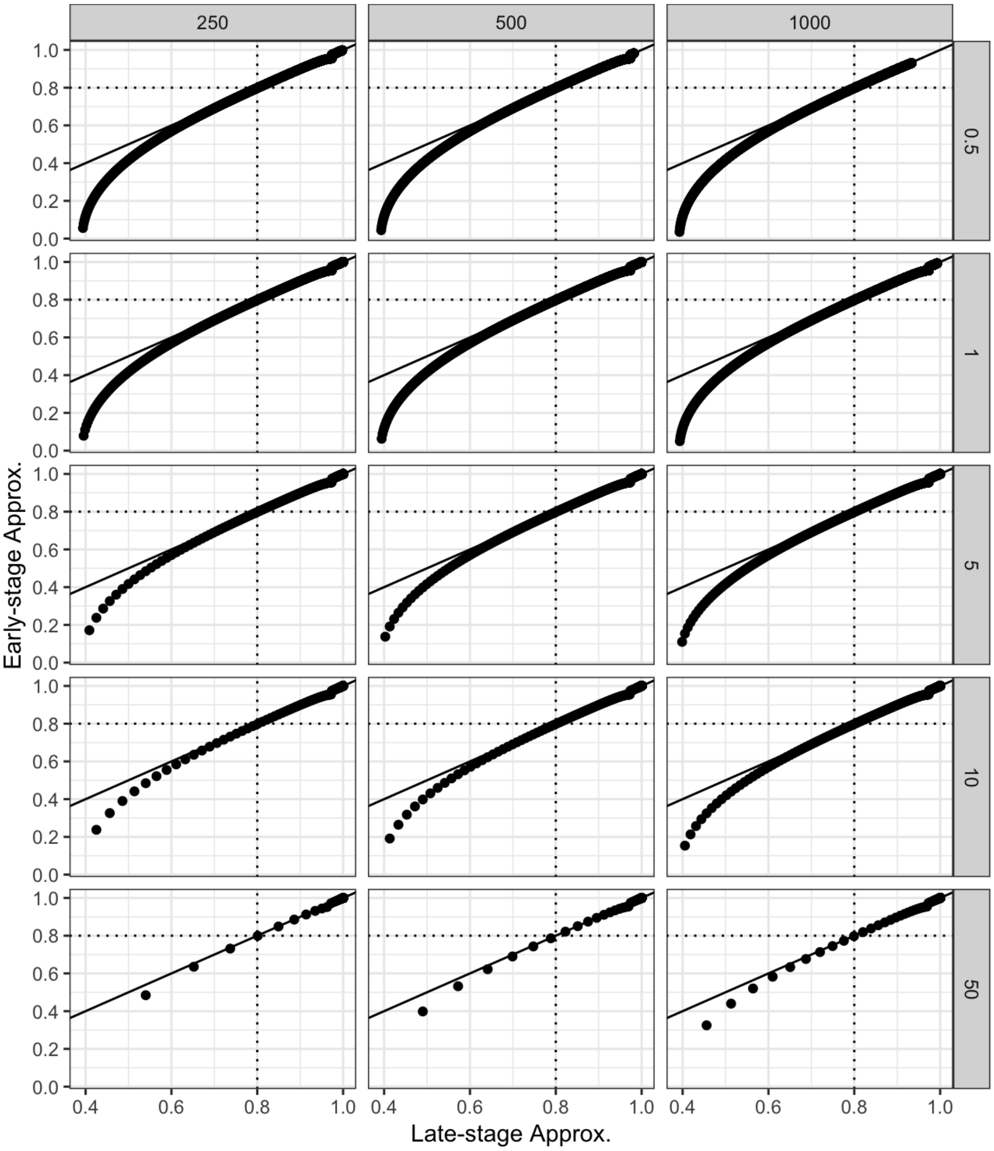

**SI Figure 13. Determination of Fickian diffusion equation approximation cutoff.** Predicted cumulative drug release fraction using the fractional release model based on early- (y-axis) and late-stage (x-axis) approximations for a sphere varying volumes (columns, $\mu$L) and diffusion coefficients (columns, $\mu$m2/s). The dotted lines at 0.8 depict the cumulative fraction where the approximations are visually similar.

**Supplementary Materials References**
